# Genetic evidence reveals the indispensable role of the *rseC* gene for autotrophy and the importance of a functional electron balance for nitrate reduction in *Clostridium ljungdahlii*

**DOI:** 10.1101/2021.08.03.455012

**Authors:** Christian-Marco Klask, Benedikt Jäger, Largus T. Angenent, Bastian Molitor

## Abstract

For *Clostridium ljungdahlii*, the RNF complex plays a key role for energy conversion from gaseous substrates such as hydrogen and carbon dioxide. In a previous study, a disruption of RNF-complex genes led to the loss of autotrophy, while heterotrophy was still possible *via* glycolysis. Furthermore, it was shown that the energy limitation during autotrophy could be lifted by nitrate supplementation, which resulted in an elevated cellular growth and ATP yield. Here, we used CRISPR-Cas12a to delete: 1) the RNF complex-encoding gene cluster *rnfCDGEAB*; 2) the putative RNF regulator gene *rseC*; and 3) a gene cluster that encodes for a putative nitrate reductase. The deletion of either *rnfCDGEAB* or *rseC* resulted in a complete loss of autotrophy, which could be restored by plasmid-based complementation of the deleted genes. We observed a transcriptional repression of the RNF-gene cluster in the *rseC*-deletion strain during autotrophy and investigated the distribution of the *rseC* gene among acetogenic bacteria. To examine nitrate reduction and its connection to the RNF complex, we compared autotrophic and heterotrophic growth of our three deletion strains with either ammonium or nitrate. The *rnfCDGEAB*- and *rseC*-deletion strains failed to reduce nitrate as a metabolic activity in non-growing cultures during autotrophy but not during heterotrophy. In contrast, the nitrate reductase deletion strain was able to grow in all tested conditions but lost the ability to reduce nitrate. Our findings highlight the important role of the *rseC* gene for autotrophy and contribute to understand the connection of nitrate reduction to energy metabolism.

**Significance Statement:** Acetogenic bacteria are widely known for their ability to convert gaseous substrates, such as hydrogen, carbon dioxide, and carbon monoxide, into short-chain fatty acids and alcohols, which can be utilized as sustainable platform chemicals and fuels. However, acetogenic bacteria conserve energy at the thermodynamic limit of life during autotrophy, and thus the production of more complex and energy-dense chemicals is limited due to low ATP yields. Therefore, it is key to decipher the interplay of the electron balancing reactions to understand and optimize the acetogenic metabolism. Recent findings with alternative electron acceptors that accelerated the cellular growth and ATP yield during autotrophy, such as nitrate, provide an opportunity to overcome energetic barriers in the acetogenic metabolism. The interrogation of the nitrate metabolism and the interplay between nitrate reduction and energy conservation in *C. ljungdahlii*, will contribute to fine-tuning of the acetogenic metabolism for biotechnological applications.

## Introduction

Acetogenic bacteria (*i.e.*, acetogens), such as *Clostridium ljungdahlii*, maintain autotrophic growth with mixtures of the gaseous substrates carbon dioxide, carbon monoxide, and hydrogen as carbon and energy sources (1, 2). The pathway that allows carbon fixation for autotrophic growth in acetogens is the Wood-Ljungdahl pathway (3, 4). Overall, the Wood-Ljungdahl pathway is considered the most energy-efficient pathway for carbon fixation that exists in nature (5, 6). In the Wood-Ljungdahl pathway, two molecules of carbon dioxide are reduced to one carbonyl group and one methyl group, which are then combined with coenzyme A to the central metabolite acetyl-coenzyme A (7). The electrons for these reductions can be derived from the oxidation of hydrogen or carbon monoxide, while carbon monoxide can also enter the pathway directly to provide the carbonyl group (8). For carbon fixation, acetyl-coenzyme A is channeled into the anabolism for cellular growth (9). For energy conservation, acetyl-coenzyme A is converted to acetate, which generates cellular energy by substrate level phosphorylation (10). One mole of ATP is generated per mole of acetate that is produced. However, in the first step of the pathway, after carbon dioxide was reduced to formate, one mole of ATP is invested to activate the formate to formyl-tetrahydrofolate (3, 4). Thus, the energy balance of the Wood-Ljungdahl pathway alone is net zero (10). All required cellular energy for the anabolism of the microbes during autotrophy is generated *via* membrane-coupled phosphorylation (2). In *C. ljungdahlii*, the membrane-bound transhydrogenase *Rhodobacter* nitrogen fixation (RNF) complex (11, 12) utilizes two electrons from the oxidation of reduced ferredoxin to reduce NAD^+^ to NADH, while simultaneously one proton is translocated across the membrane (10, 13). A proton-dependent F_1_F_O_ ATPase then consumes the chemiosmotic proton gradient to generate ATP (14, 15). In the presence of carbon dioxide and hydrogen, theoretically, *C. ljungdahlii* can generate a maximum of 0.63 moles ATP per mole acetate for the anabolism *via* membrane-coupled phosphorylation. Thus, the conservation of cellular energy during autotrophy occurs at the thermodynamic limit of life (10).

For *C. ljungdahlii*, the RNF complex is encoded by the RNF-gene cluster *rnfCDGEAB*. Although the RNF complex plays an essential role for energy conservation during autotrophy in *C. ljungdahlii* (13), fundamental knowledge about the regulation and gene expression control of the encoding RNF-gene cluster is missing. Transcriptome studies with *C. ljungdahlii* revealed that the RNF complex is under strict gene expression control and strongly induced during autotrophy (15, 16). The regulatory mechanisms behind this remain unknown. However, the small gene *rseC*, which is located directly upstream of *rnfC* in *C. ljungdahlii*, is also highly expressed during autotrophy and follows the expression profile of *rnfC* (15). The gene *rseC* is annotated to contain the conserved protein domain family RseC_MucC (pfam04246) (14). The domain family RseC_MucC is found in positive transcriptional regulators in other microbes. The one representative, RseC, was found to be involved in the oxidative stress response in *Escherichia coli* (17–19), and in thiamine synthesis in *Salmonella typhimurium* (20). The other representative, MucC, was found to be involved in the regulation of the alginate formation of *Azotobacter vinelandii* (21) and *Pseudomonas aeruginosa* (22). Others identified a transcription start site for *C. ljungdahlii*, which is located upstream of the *rseC* gene, and a putative terminator sequence, which is located between *rseC* and *rnfC* were identified. This indicates that *rseC* is expressed as an individual transcript apart from the RNF-gene cluster transcripts (15). Altogether, this leads to the assumption that the *rseC* gene product is closely linked to the RNF complex, and could be important for the regulation of autotrophy in *C. ljungdahlii*.

While autotrophy in acetogens results in low cellular energy yields, Emerson *et al.* (23) reported that *C. ljungdahlii* is able to couple the reduction of nitrate to the generation of ATP during growth with carbon dioxide and hydrogen. This relieved the energy limitation during autotrophy and resulted in a significantly higher biomass yield (23). We confirmed this in a bioreactor study, and biomass yields were considerably higher with nitrate, but resulted in stochastic crashes of the continuous bioreactor cultures (24). Emerson *et al.* (23) proposed that electrons, which are required for nitrate reduction, are provided by NADH. One route to regenerate NADH is by the RNF complex where reduced ferredoxin is consumed (11), which would link nitrate reduction to the energy metabolism. It was assumed that nitrate reduction is accelerating the RNF-complex activity and more protons are translocated across the membrane, which can be used by the F_1_F_O_ ATPase for the generation of ATP (23). This way, the co-utilization of carbon dioxide and nitrate with hydrogen was suggested to yield up to 1.5 ATP through the concerted action of the RNF complex and the ATPase (23). This would be a 2.4-fold increase in ATP yield compared to the ATP yield with carbon dioxide and hydrogen alone (10).

To investigate the autotrophy in *C. ljungdahlii* with respect to regulatory aspects and the interplay with nitrate reduction, we addressed three main questions: *1)* Is the *rseC* gene involved in the regulation of the RNF-gene cluster?; *2)* Is nitrate reduction dependent on a functional RNF complex?; and *3)* Is nitrate reduction abolished by the deletion of the nitrate reductase that is annotated in the genome of *C. ljungdahlii*?

## Results

### A full deletion of the RNF complex confirmed its indispensable role for autotrophy in *C. ljungdahlii*

We first attempted to generate a full deletion of the RNF-gene cluster to further investigate autotrophy in *C. ljungdahlii*. Others had demonstrated that a mutant strain of *C. ljungdahlii*, for which the *rnfAB* genes were disrupted with an antibiotic resistance cassette, had lost the ability to grow during autotrophy (13). However, this genome modification was not stable, and the wild-type genotype was restored during the cultivation time of the experiments (13). In addition, it was demonstrated recently that a full RNF-gene cluster deletion led to the loss of autotrophic growth in the acetogen *Acetobacterium woodii* (25). Here, we achieved a full deletion of the RNF-gene cluster in *C. ljungdahlii* with a clustered regularly interspaced short palindromic repeats (CRISPR)-associated protein 12a (CRISPR-Cas12a) system, which we implemented and used to generate all deletion strains in this study (Fig. 1A, **Supplementary Text S1A**).

**Fig. 1.**
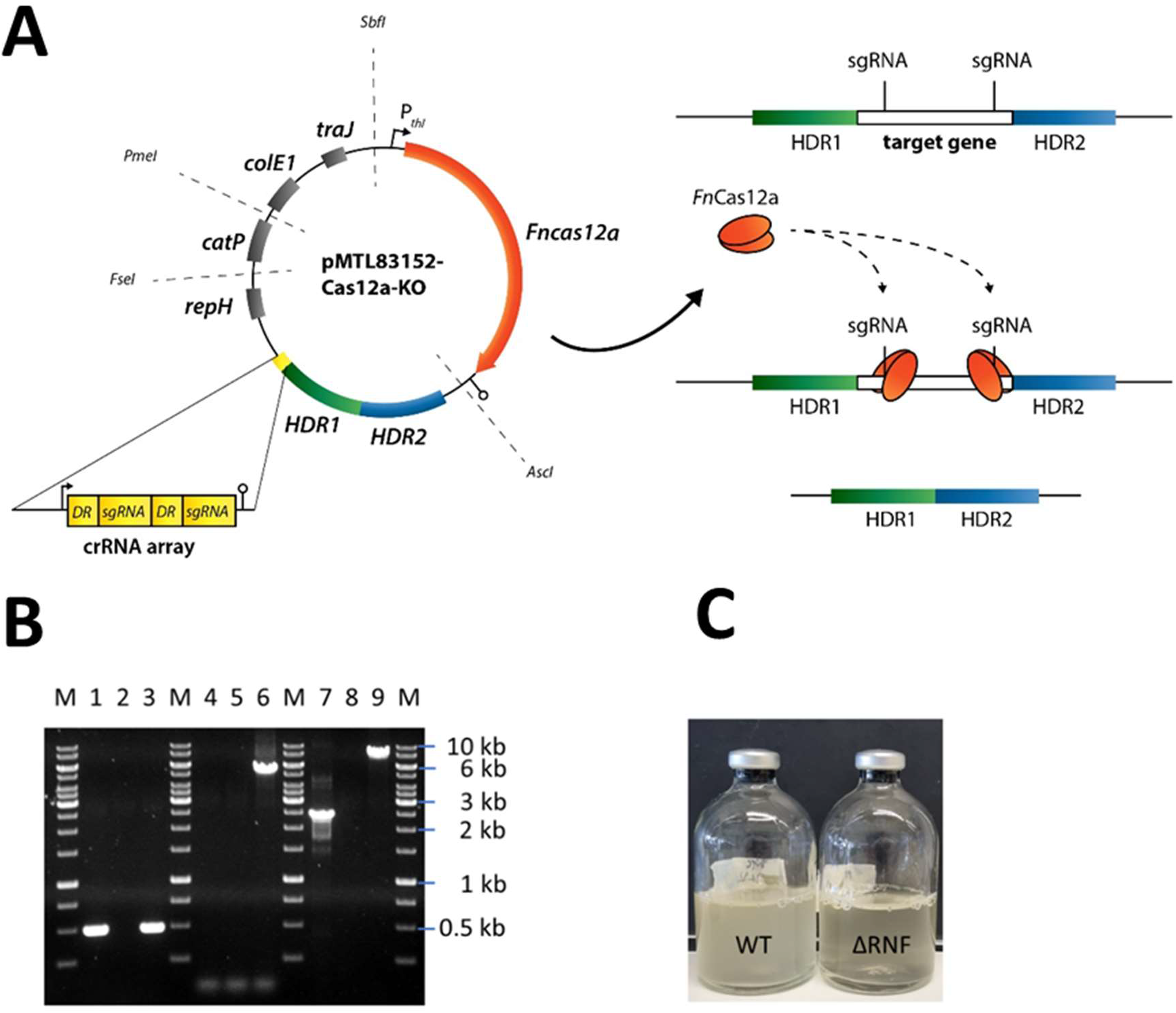
CRISPR-Cas12a-mediated rnfCDGEAB gene cluster deletion in *C. ljungdahlii*. **A,** modular CRISPR-Cas12a system established in the pMTL80000 shuttle-vector system (37). The final CRISPR-Cas12a plasmid for deletion of *rnfCDGEAB* contained the *Fncas12a* gene, homology-directed repair arms (HDRs), and a specific crRNA array comprising two directed repeats (DRs) and two sgRNA, which targeted the *rnfC* and *rnfB* genes. B, agarose gel with PCR-samples for the *fdhA* fragment (WT: 501 bp, deletion strain: 501 bp), *rnfCDGEAB* fragment (WT: 5047 bp, deletion strain: no fragment), and for a fragment that was amplified with primers that bind ∼1250 bp upstream and downstream of the *rnfCDGEAB* gene cluster locus (WT: 7550 bp, deletion strain: 2503 bp). DNA-template: gDNA of *C. ljungdahlii* ΔRNF (lane 1, 4, and 7); gDNA of *C. ljungdahlii* WT (lane 3, 6, and 9); and water (lane 2, 5, 8). M: Generuler^TM^ 1 kb DNA ladder. **C**, growth of the wild type (WT) and reduced growth of the deletion strain (ΔRNF) with fructose in PETC medium. HDR1/2, homology-directed repair arm flanking the targeted gene; crRNA array, sequence containing FnCas12a-specific DRs and sgRNAs; sgRNA, guide RNA; *repH*, Gram-positive origin of replication; *catP*, antibiotic resistant cassette against chloramphenicol/thiamphenicol; *colE1*, Gram-negative origin of replication; *traJ*, conjugation gene; **P***_thl_*, promoter sequence of the thiolase gene in *Clostridium acetobutylicum*; *Asc*I, *Fse*I, *Pme*I, and *Sbf*I are unique-cutting restriction sites, which were preserved during the cloning to maintain the modular functionality of the plasmid backbone.

After successfully generating the RNF-gene cluster deletion strain (*C. ljungdahlii* ΔRNF) and confirming the identity of this strain (Fig. 1B, **Supplementary Text S1B**), we compared the growth of *C. ljungdahlii* wild-type (WT) to the growth of *C. ljungdahlii* ΔRNF. We performed growth experiments with carbon dioxide and hydrogen (autotrophy) and with fructose (heterotrophy), while we added equimolar amounts of either ammonium or nitrate as nitrogen source to the medium for both autotrophy and heterotrophy (**Materials and Methods**, Fig. 2, **Supplementary Fig. S1**). We had added a small amount of yeast extract (0.1 weight-%) in all cultivation conditions (**Material and Methods**). As expected, we observed growth for *C. ljungdahlii* WT in all growth experiments (Fig. 2A, **Supplementary Fig. S1A**). However, the nitrogen source had a distinct influence on the growth rate, final OD_600_, fermentation product spectrum, and pH (*Table 1*, Fig. 2B, **Supplementary Text S1C, Supplementary Fig. S1B**). We found that nitrate reduction occurred rapidly in our growth experiments (Fig. 2F, **Supplementary Fig. S1F**). *C. ljungdahlii* WT utilized all provided nitrate within 53 h of cultivation with carbon dioxide and hydrogen (Fig. 2F) and within 47 h of cultivation with fructose (**Supplementary Fig. S1F**). The ammonium concentrations increased concomitant with decreasing nitrate concentrations when nitrate was provided in the medium (Fig. 2E, **Supplementary Fig. S1E**). Noteworthy, we also observed an increase in the ammonium concentration when ammonium was provided as the nitrogen source during autotrophy (Fig. 2F). We did not measure any nitrite as an intermediate of the nitrate reduction pathway (discussed below).

**Fig. 2.**
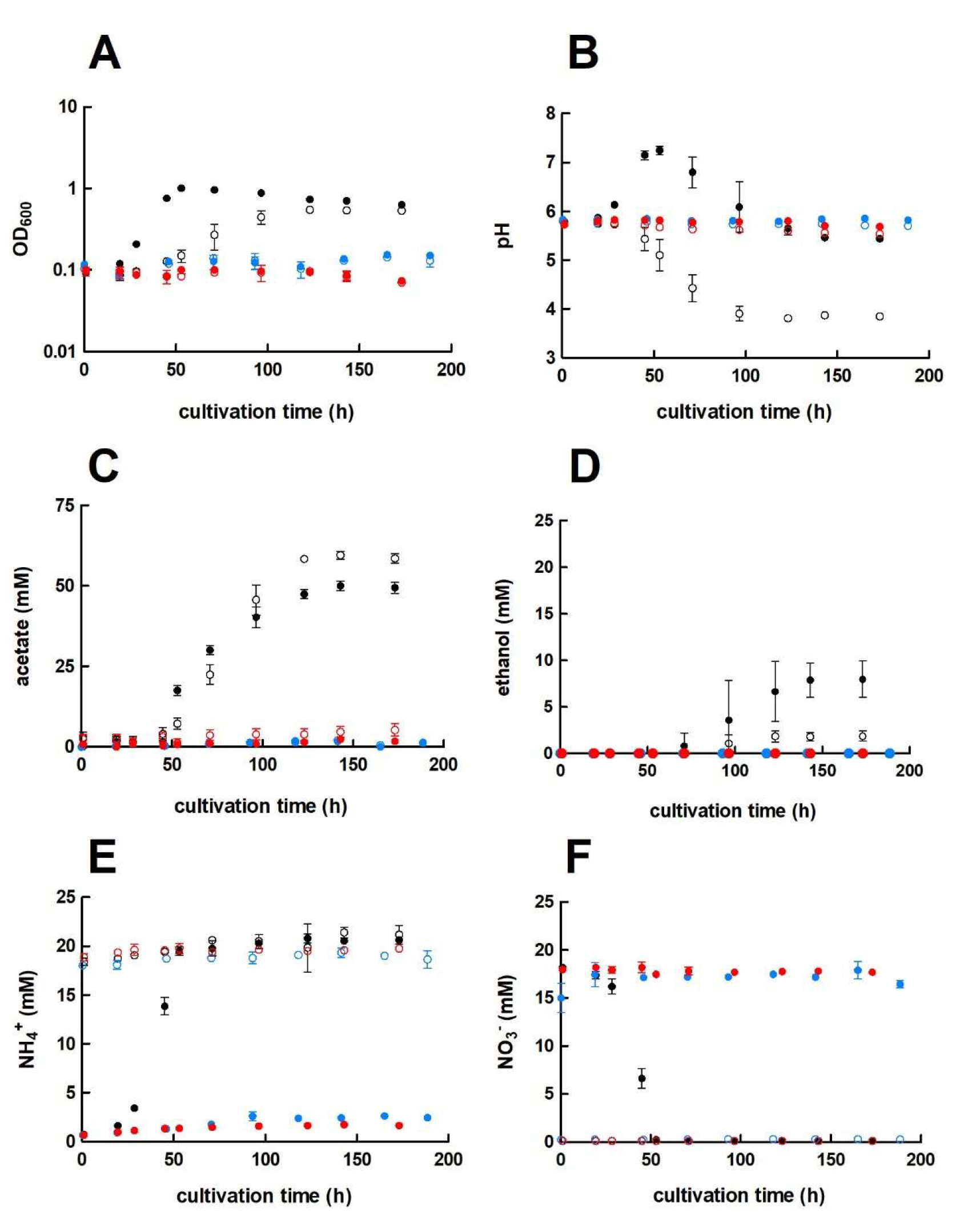
Cultivation of *C. ljungdahlii* WT, *C. ljungdahlii* ΔRNF, and C. ljungdahlii ΔrseC in nitrate-or ammonium-containing medium with H_2_ and CO_2_. Cultures of *C. ljungdahlii* strain WT (●, ○), ΔRNF (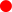, 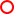), and Δ*rseC* (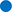, 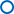) were grown in 100 mL PETC medium in 1 L bottles at 37°C and 150 rpm. The headspace consisted of H_2_ and CO_2_ (80/20 vol-%) and was set to 0.5 bar overpressure. The medium contained either 18.7 mM nitrate (NO_3_^-^) (filled circles) or 18.7 mM ammonium (NH_4_^+^) (open circles) as nitrogen source. The cultivation times were 173 h for cultures of *C. ljungdahlii* WT and *C. ljungdahlii* ΔRNF and 186 h for cultures of *C. ljungdahlii* Δ*rseC*. All cultures were grown in biological triplicates, data is given as mean values, with error bars indicating the standard deviation. **A**, growth; **B**, pH-behavior; **C**, acetate concentrations; **D**, ethanol concentration; **E**, ammonium concentration; and **F**, nitrate concentrations. WT, wild type; ΔRNF, RNF-gene cluster deletion; Δ*rseC*, *rseC* gene deletion.

**Table 1.** Performance of all tested *C. ljungdahlii* strains in autotrophic batch cultivation experiments. Cultures were grown with carbon dioxide and hydrogen (autotrophy) in PETC medium, which contained either ammonium or nitrate as nitrogen source. A gas atmosphere of H_2_/CO_2_ (80/20 vol-%) with 0.5 bar overpressure was applied. Growth was not detected for any culture of *C. ljungdahlii* ΔRNF or *C. ljungdahlii* Δ*rseC*. Data is represented as mean values from biological triplicates ± standard deviation. WT, *C. ljungdahlii* wild type; ΔRNF, *C. ljungdahlii* with deleted *rnfCDGEAB* gene cluster; Δ*rseC*, *C. ljungdahlii* with deleted *rseC* gene; and Δ*nar, C. ljungdahlii* with deleted nitrate reductase gene cluster. CO_2_, carbon dioxide; and H_2_, hydrogen.

| strain | nitrogen source | growth rate (μ in h)<sup>a</sup> | maximum OD<sub>600</sub> value | maximum acetate concentration (mM) | maximum ethanol concentration (mM) |
| --- | --- | --- | --- | --- | --- |
| WT | ammonium | 0.024±0.002 | 0.56±0.01 | 59.5±1.8 | 1.9±0.4 |
| WT | nitrate | 0.072±0.004 | 1.00±0.06 | 50.1±2.1 | 8.0±1.6 |
| ΔRNF | ammonium | - | - | 5.7±3.0 | n.d.<sup>b</sup> |
| ΔRNF | nitrate | - | - | 2.3±1.1 | n.d.<sup>b</sup> |
| ΔrseC | ammonium | - | - | 2.0±0.5 | n.d.<sup>b</sup> |
| ΔrseC | nitrate | - | - | 1.9±0.1 | n.d.<sup>b</sup> |
| Δnar | ammonium | 0.018±0.001 | 0.44±0.01 | 44.8±0.2 | 3.3±0.2 |
| Δnar | nitrate | 0.017±0.003 | 0.44±0.01 | 41.9±1.9 | 2.9±0.4 |

<sup>a</sup> μ values were calculated based on the individual OD<sub>600</sub> values of each triplicate in the exponential growth phase.
<sup>b</sup> n.d., not detectable.

In contrast, the *C. ljungdahlii* ΔRNF strain was unable to grow with carbon dioxide and hydrogen regardless of the nitrogen source (Fig. 2A). We did not observe a pH decrease, and also not an accumulation of ethanol as a metabolic activity of non-growing cultures of *C. ljungdahlii* ΔRNF, however, some minor amounts of acetate were detected (Fig. 2B, 2C, 2D, Table 1). Furthermore, nitrate reduction as a metabolic activity of non-growing cultures was not detectable in *C. ljungdahlii* ΔRNF with carbon dioxide and hydrogen (Fig. 2F). Thus, we confirmed the essential role of the RNF complex for autotrophy in *C. ljungdahlii*.

### The deletion of the RNF complex influenced nitrate reduction during heterotrophy

For the *C. ljungdahlii* ΔRNF strain, heterotrophic growth with fructose was still possible but notably reduced (Fig. 1C, **Supplementary Fig. S1A**). The growth rates of *C. ljungdahlii* ΔRNF were significantly reduced by 34% (0.052 h^-1^, *P* ≤ 0.001) and by 42% (0.042 h^-1^, *P* ≤ 0.001) with ammonium and nitrate, respectively, when compared to *C. ljungdahlii* WT (Table 1, **Supplementary Table S1**). The observed maximum OD_600_ values were also significantly reduced by 53% (*P* ≤ 0.001) and 56% (*P* ≤ 0.001) for *C. ljungdahlii* ΔRNF, respectively (Table 1, **Supplementary Table S1A**). In addition, the maximum acetate concentrations in the deletion strain were significantly reduced by 32% (*P* ≤ 0.001) with ammonium and by 42% (*P* ≤ 0.001) with nitrate compared to the maximum acetate concentration in the wild type. The maximum ethanol concentration was significantly reduced by 41% (*P* ≤ 0.001) with fructose and ammonium, while ethanol was not produced at all by *C. ljungdahlii* ΔRNF during growth with fructose and nitrate (Table 1, **Supplementary Fig. S1C, S1D**). During heterotrophy, *C. ljungdahlii* ΔRNF was able to utilize nitrate but considerably slower when compared to *C. ljungdahlii* WT (**Supplementary Fig. S1F**). At the end of the cultivation, cultures of *C. ljungdahlii* ΔRNF had only consumed 49% of the provided nitrate (**Supplementary Fig. S1F**). Overall, we observed a halt in growth and metabolic activity for cultures of *C. ljungdahlii* ΔRNF with fructose after 47 h of cultivation in nitrate-containing medium and after 56 h of cultivation in ammonium-containing medium (**Supplementary Fig. S1**). Fructose concentrations at the end of the cultivation remained at a concentration of 8.0-9.7 mM, which is still 30-35% of the initially provided concentration (**Supplementary Fig. S1G**). The pH did not increase during heterotrophy with nitrate in *C. ljungdahlii* ΔRNF, but instead slowly decreased until the end of the cultivation (**Supplementary Fig. S1B**). Notably, the final pH for heterotrophic cultures of *C. ljungdahlii* ΔRNF with nitrate was still higher compared to cultures with ammonium (**Supplementary Fig. S1B**). For none of the culture samples with *C. ljungdahlii* ΔRNF during heterotrophy, decreasing ammonium concentrations were observed, even when ammonium was provided as the nitrogen source (**Supplementary Fig. S1E**). Overall, we confirmed that the RNF complex plays a pivotal role for the distribution of electrons in the metabolism of *C. ljungdahlii* during heterotrophy, but that it was not essential in these conditions.

### The *rseC* gene is essential for autotrophy in *C. ljungdahlii*

After we confirmed the indispensable role of the RNF complex for autotrophy and the influence on nitrate reduction during heterotrophy, we investigated the role of the small putative regulator gene *rseC* (CLJU_c11350), which is located directly upstream of the *rnfCDGEAB* gene cluster. A transcriptomic study with *C. ljungdahlii* had revealed that *rseC* is expressed in a similar pattern compared to *rnfC* and is highly expressed during autotrophy (15). We applied our CRISPR-Cas12a system to delete the *rseC* gene from the genome (**Supplementary Fig. S2A**). Next, we performed growth experiments with the generated *C. ljungdahlii* Δ*rseC* strain under the same conditions as for the *C. ljungdahlii* WT and ΔRNF strains. Cultures of *C. ljungdahlii* Δ*rseC* did not grow with carbon dioxide and hydrogen, neither with ammonium nor with nitrate, during a total cultivation time of 189 h (Fig. 2). Non-growing cultures for this strain did not accumulate notable concentrations of acetate or ethanol during the cultivation time (Fig. 2C, 2D). Furthermore, we did not observe nitrate reduction or a remarkable change in pH as a metabolic activity of non-growing cultures for this strain during autotrophy (Fig. 2B, 2E, 2F).

Heterotrophic growth of *C. ljungdahlii* Δ*rseC* was possible, and in contrast to *C. ljungdahlii* ΔRNF, the impact was less pronounced for growth with ammonium but limited to some extent with nitrate (**Supplementary Fig. S1A**). Heterotrophic growth rates were increased by 6% (0.084 h^-1^, *P* = 0.08) with ammonium and significantly reduced by 34% (0.048 h^-1^, *P* ≤ 0.001) with nitrate as nitrogen source, respectively, when compared to *C. ljungdahlii* WT under the same conditions (**Supplementary Table 1**). The maximum observed OD_600_ values for *C. ljungdahlii* Δ*rseC* were 1.90±0.15 for ammonium and 1.58±0.03 for nitrate cultures, which is a reduction of 24% (*P* = 0.05) and a significant reduction of 30% (*P* ≤ 0.001) compared to the wild type. The maximum acetate concentrations of *C. ljungdahlii* Δ*rseC* were similar to those observed for *C. ljungdahlii* WT, while the maximum ethanol concentrations were significantly reduced by 29% (*P* ≤ 0.001) for ammonium cultures and by 42% (*P* ≤ 0.001) for nitrate cultures instead (**Supplementary Table 1**, **Supplementary Fig. S1C, S1D**). Nitrate reduction was not restricted during heterotrophy in *C. ljungdahlii* Δ*rseC* (**Supplementary Fig. S1E, S1F**). Indeed, we observed a rapid utilization of all supplied nitrate within 60 h of cultivation, which is similar to the observations that we had made for *C. ljungdahlii* WT (**Supplementary Fig. S1F**). Thus, *rseC* seems to be involved in positively regulating the expression of the RNF-gene cluster during autotrophy, but not during heterotrophy. However, the exact impact on gene expression of the RNF-gene cluster cannot be deduced from these findings.

### Plasmid-based complementation relieved the phenotypes of the *C. ljungdahlii* ΔRNF and Δ*rseC* strains

After we had characterized the *C. ljungdahlii* ΔRNF and *C. ljungdahlii* Δ*rseC* strains, we questioned whether the wild-type phenotype, particularly with respect to autotrophy, can be restored by plasmid-based gene complementation. Therefore, we generated the plasmid-carrying strains *C. ljungdahlii* ΔRNF pMTL83151_P_nat__*rnfCDGEAB* and *C. ljungdahlii* Δ*rseC* pMTL83152_*rseC*. The plasmids encode the RNF-gene cluster under the control of the native promoter region upstream of the *rnfC* gene from the genome (P_nat_) in pMTL83151_P_nat__*rnfCDGEAB* and the *rseC* gene under the control of the constitutive thiolase promoter (P*_thl_*) in pMTL83152_*rseC*, respectively. We investigated the complementation strains in ammonium-containing medium with carbon dioxide and hydrogen for growth (Fig. 3). Indeed, the plasmid-based expression of the deleted genes relieved the phenotype and enabled autotrophy with carbon dioxide and hydrogen for both strains (Table 2, Fig. 3). The control strains that carried an empty plasmid failed to grow autotrophically, as we had already observed for the non-complemented deletion strains.

**Fig. 3.**
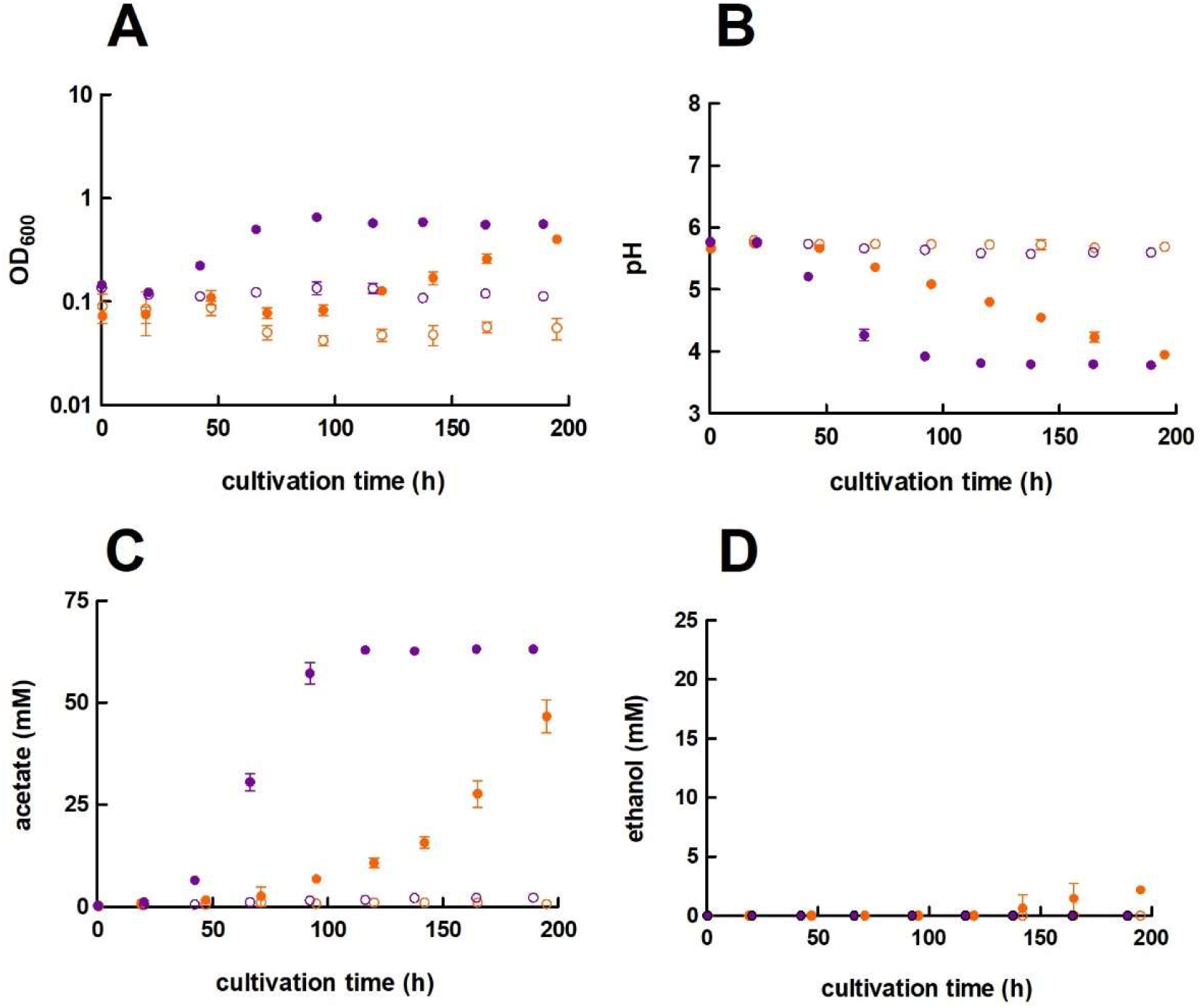
Growth and pH behavior of plasmid-based complementation of *C. ljungdahlii* ΔRNF and *C. ljungdahlii* Δ*rseC* with H_2_ and CO_2_. Cultures were grown in 100 mL PETC medium in 1 L bottles at 37°C and 150 rpm for 195 h and 189 h, respectively. The headspace consisted of H_2_ and CO_2_ (80/20 vol-%) and was set to 0.5 bar overpressure. Only 18.7 mM ammonium (NH_4_) but no nitrate was added to the medium. All cultures were grown in biological triplicates, data is given as mean values, with error bars indicating the standard deviation. **A**, growth and **B**, pH-behavior of *C. ljungdahlii* ΔRNF strains. **C**, growth and **D**, pH-behavior of *C. ljungdahlii* Δ*rseC* strains. 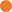 *C. ljungdahlii* ΔRNF pMTL83151_P_nat_*_rnfCDGEAB*; 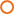 *C. ljungdahlii* ΔRNF pMTL83151; 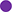 *C. ljungdahlii* Δ*rseC* pMTL83152_*rseC*; and 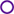 *C. ljungdahlii* Δ*rseC* pMTL83152. ΔRNF, rnfCDGEAB gene cluster deletion; Δ*rseC*, deletion of *rseC*; P_nat_, native promoter sequence upstream of *rnfC*; P*_thl_*, promoter of the thiolase gene in *C. acetobutylicum*; rpm, revolutions per minute; CO_2_, carbon dioxide; and H_2_, hydrogen.

**Table 2.**
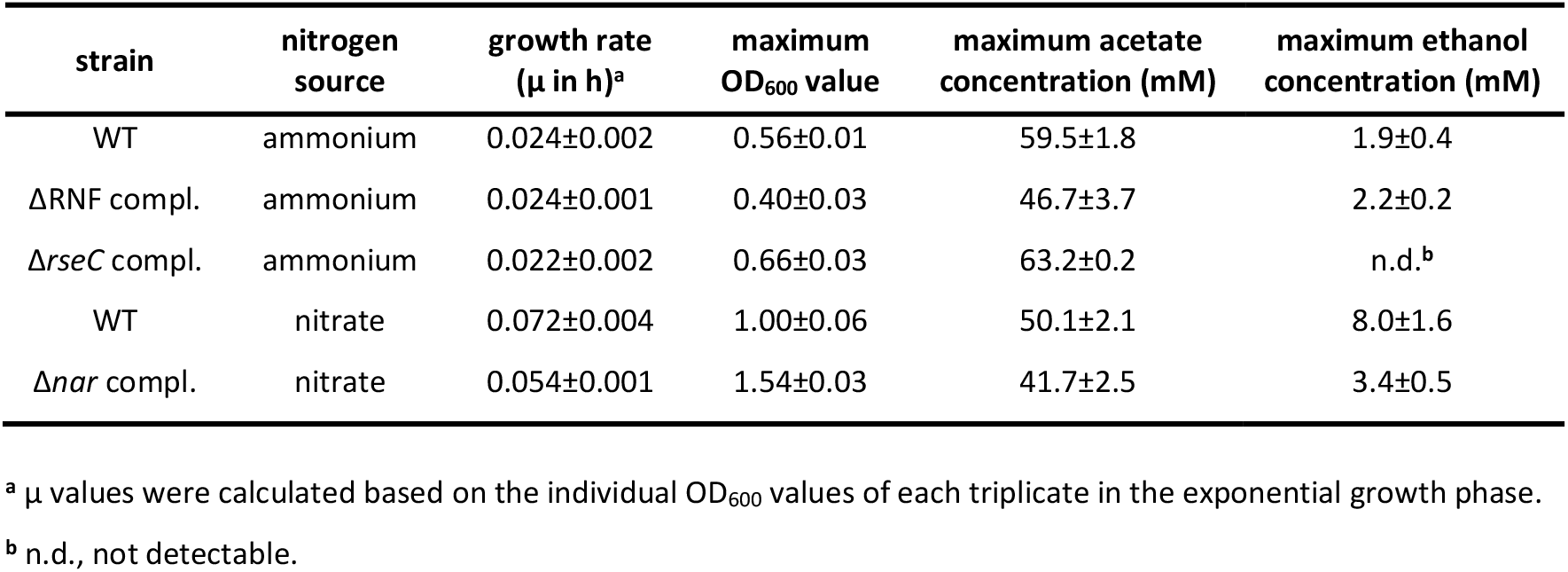
Performance of the plasmid-based complemented deletion strains of *C. ljungdahlii* in autotrophic batch cultivation experiments. Cultures were grown with carbon dioxide and hydrogen (autotrophy) in PETC medium, which contained either ammonium or nitrate as nitrogen source. A gas atmosphere of H_2_/CO_2_ (80/20 vol-%) with 0.5 bar overpressure was applied. Data is represented as mean values from biological triplicates ± standard deviation. The WT data from Table 1 are shown again for comparison. WT, wild type; ΔRNF, deletion of the *rnfCDGEAB* gene cluster; Δ*rseC*, deletion of the *rseC* gene; Δ*nar*, deletion of the nitrate reductase gene cluster; ΔRNF compl., complementation strain *C. ljungdahlii* pMTL83151_P_nat__*rnfCDGEAB*; Δ*rseC* compl., complementation strain *C. ljungdahlii* pMTL83152_*rseC*; and Δ*nar* compl., complementation strain *C. ljungdahlii* pMTL83152_*nar*.

However, *C. ljungdahlii* ΔRNF pMTL83151_P_nat__*rnfCDGEAB* reached only 71% (*P* = 0.02) of the maximum OD_600_ with ammonium when compared to the wild type, which is significantly less (Table 2). Furthermore, the complemented strain had a prolonged *lag* phase of 71 h (Fig. 3A). The pH decrease occurred slower compared to the wild type (Fig. 3B). The *C. ljungdahlii* ΔRNF pMTL83151_P_nat__*rnfCDGEAB* strain reached a maximum acetate concentration of 46.7±3.4 mM, which is a significant reduction of 22% (*P* = 0.009) when compared to the wild type (Table 2, Fig. 3C). The maximum ethanol concentration was similar in comparison to the wild type (Fig. 3D). In contrast, the *C. ljungdahlii* Δ*rseC* pMTL83152_*rseC* strain reached a maximum OD_600_ of 0.66±0.03, which is a significant increase of 17% (*P* = 0.02) compared to the wild type (Table 2, Fig. 3A). Instead of a prolonged *lag* phase, we observed a shortened *lag* phase for this strain when compared to the wild type (Fig. 1A, 3A). Notably, the medium for the complementation experiments always contained antibiotics, which generally caused a slightly negative impact on growth of plasmid-carrying *C. ljungdahlii* strains such as in the *C. ljungdahlii* ΔRNF pMTL83151_P_nat__*rnfCDGEAB* strain. In contrast, this was not the case for the *C. ljungdahlii* Δ*rseC* pMTL83152_*rseC* strain. The complemented strain reached a maximum acetate concentration of 63.2±0.2 mM (Fig. 3C), which is a significant increase of 6% (*P* = 0.04) when compared to the wild type (Table 2, Fig. 2). However, this strain did not produce any detectable ethanol during the cultivation (Table 2, Fig. 3D). Furthermore, the pH value did not show any notable change, when compared to the wild type (Fig. 3B).

### Plasmid-based overexpression of the *rseC* gene enhanced autotrophic growth

We observed a growth stimulating effect in the *C. ljungdahlii* Δ*rseC* pMTL83152_*rseC* strain. To investigate whether overexpression of *rseC* in the wild-type strain increases autotrophic growth further, we generated the *C. ljungdahlii* pMTL83152_*rseC* strain. This strain carries the complementation plasmid with the constitutive P*_thl_* promoter in the wild-type background. During autotrophy with carbon dioxide and hydrogen in ammonium-containing medium, the *C. ljungdahlii* pMTL83152_*rseC* strain had a shortened *lag* phase and a 13.2% faster but not significantly increased growth rate (0.21 h^-1^, *P* = 0.2) compared to the wild type (Fig. 2A, **Supplementary Fig. S3A**). In addition, this overexpression strain reached similar maximum OD_600_ values (**Supplementary Fig. S3A**). The maximum acetate concentration was significantly reduced by 22% (*P* ≤ 0.001), and ethanol was not produced (**Supplementary Fig. S3C, S3D**).

We also attempted to generate a plasmid that carries the *rnfCDGEAB* gene cluster under the control of a constitutive promoter. However, any attempts to generate a fusion of the constitutive promoter P*_thl_* with the *rnfCDGEAB* gene cluster failed already during the cloning steps in *E. coli*. Thus, for the expression of *rnfCDGEAB* in the wild type, we also used the native P_nat_ promoter sequence, which most likely is under the same expression control as the genomic copy of the RNF-gene cluster. Not surprisingly, the cultivation of *C. ljungdahlii* pMTL83151_P_nat__*rnfCDGEAB* did not show any notable impact on growth and product formation when compared to the control strain that carried an empty plasmid (**Supplementary Fig. S3**).

### The gene expression profiles of rnf genes and the *rseC* gene in the deletion strains revealed regulatory effects

We had found that autotrophy was abolished in the *rseC* deletion strain, while heterotrophy was not impacted. Thus, we further investigated the activating or repressing function on the gene expression of the RNF-gene cluster by RseC. For this, we performed qRT-PCR analyses to investigate the individual expression profiles of the genes *rnfC, rnfD, rnfG, rnfE, rnfA, rnfB*, and *rseC* in the *C. ljungdahlii* Δ*rseC* strain. We included the *C. ljungdahlii* ΔRNF and *C. ljungdahlii* WT strains as controls (*Materials and Methods*). We analyzed samples after 3 h and 20 h of cultivation time to investigate the transcriptomic response after inoculating the autotrophic and heterotrophic main cultures from heterotrophic pre-cultures. During the cultivation of the six main cultures (three strains, two conditions), *C. ljungdahlii* WT grew during autotrophy and heterotrophy, while *C. ljungdahlii* ΔRNF and *C. ljungdahlii* Δ*rseC* only grew during heterotrophy.

The qRT-PCR results in this paragraph are given as log_2_ (fold change in gene expression), where a value of ≤ −1 (0.5-fold) refers to a significant downregulation, and a value of ≥ +1 (2-fold) refers to a significant upregulation (Fig. 4). We did not measure any expression signals for any of the deleted RNF genes in the *C. ljungdahlii* ΔRNF strain and for the deleted *rseC* gene in the *C. ljungdahlii* Δ*rseC* strain. We found that all RNF-gene cluster genes were significantly downregulated (ranging from −1.8 to −4.7) in the *C. ljungdahlii* Δ*rseC* strain, when cultivating non-growing cells of this strain autotrophically with hydrogen and carbon dioxide (Fig. 4). We observed a similar pattern of downregulation for the 3-h and 20-h samples of the *C. ljungdahlii* Δ*rseC* strain (Fig. 4A and 4C). In the heterotrophic samples, all RNF-gene cluster genes, except of *rnfB*, were significantly downregulated in the 3-h samples (ranging from −1.0 to −1.8). However, after 20 h of cultivation time during heterotrophy, we observed a less pronounced but still significant downregulation of the *rnfC* gene, while all other genes were either not significantly different from the wild type (*rnfD*) or significantly upregulated (Fig. 4). In the *C. ljungdahlii* ΔRNF strain as a control, we found that *rseC* expression was significantly upregulated in the 3-h samples during autotrophy (+2.4) and during heterotrophy (+1.3) (Fig. 4B). The upregulation was less pronounced but still significant after 20 h of cultivation (Fig. 4D). For the wild type, all genes (except for *rnfE* in the 3-h sample) were significantly upregulated during autotrophy when compared to heterotrophy for the 3-h samples (ranging from +1.0 to +5.4), and for the 20 h samples (ranging from +2.8 to +3.8), respectively (**Supplementary Fig. S4**). Thus, RseC positively regulated the RNF-gene cluster during autotrophy, but not during heterotrophy.

**Fig. 4.**
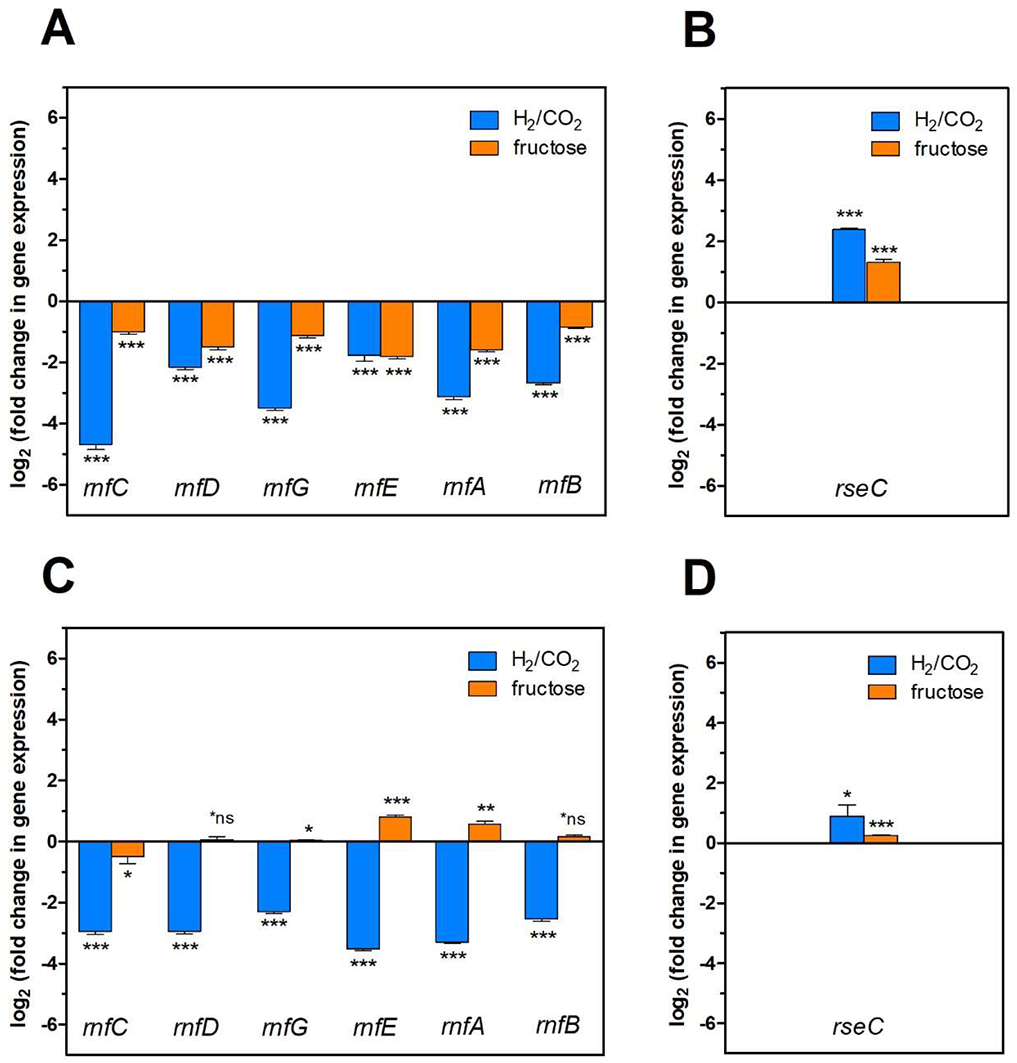
Gene expression change of the *rnfCDGEAB* cluster genes and the *rseC* gene in the ΔRNF and Δ*rseC* deletion strains. **A,** gene expression change for the genes *rnfC*, *rnfD*, *rnfG*, *rnfE*, *rnfA*, and *rnfB* in strain *C. ljungdahlii* Δ*rseC* after 3h cultivation time; **B**, gene expression change for the gene *rseC* in strain *C. ljungdahlii* ΔRNF after 3 h cultivation time; **C**, gene expression change for the genes *rnfC*, *rnfD*, *rnfG*, *rnfE*, *rnfA*, and *rnfB* in strain *C. ljungdahlii* Δ*rseC* after 20 h cultivation time; and **D**, gene expression change for the gene *rseC* in strain *C. ljungdahlii* ΔRNF after 20 h cultivation time. RNA samples were purified from cultures that were cultivated either autotrophically with hydrogen and carbon dioxide (blue bars) or heterotrophically with fructose (orange bars). cDNA was synthesized from the purified RNA samples and used as template for qRT-PCR analyses. The individual gene expression profiles of each gene was calculated using the wild-type strain as reference, which was grown under the same conditions. The *rho* gene was used as “housekeeping” gene. The fold change in gene expression was determined with the 2^-ΔΔCT^ method (43). ***, *P* ≤ 0.001; **, *P* ≤ 0.01; *, *P* ≤ 0.05; *ns, not significant (*P* > 0.05). We defined log_2_ (fc) ≤ −1 as downregulated genes and ≥ +1 as upregulated genes.

### The *rseC* gene is abundantly found among acetogens

While *rseC* was annotated as a putative transcriptional regulator, the regulatory function was not known. We had found in our cultivation experiments and qRT-PCR analyses that the *rseC* gene plays a critical role for the function of the RNF complex, and thus for autotrophy. We investigated whether *rseC* genes are also present in genomes of other microbes that possess RNF complex genes. Indeed, we found putative *rseC* genes in the genomes of *C. ljungdahlii*, *Clostridium autoethanogenum, A. woodii*, *Eubacterium limosum, Clostridium carboxidovorans, Clostridium kluyveri, R. capsulatus*, and *E. coli*. On the contrary, we did not find a putative *rseC* gene in the genome of *Moorella thermoacetica* or *Thermoanaerobacter kivui*, which possess an energy-converting hydrogenase (Ech) complex instead of an RNF complex (26). Next, we took a detailed look at the genomic location and distance to the RNF-gene cluster (Fig. 5). We noticed that the *rseC* gene was located directly upstream of the RNF complex gene cluster in *C. ljungdahlii* (CLJU_c11350), *C. autoethanogenum* (CAETHG_3225), *C. carboxidovorans* (Ccar_25725), and *C. kluyveri* (CKL_1263). The *rseC* gene in *A. woodii* (Awo_C21740) and *E. limosum* (B2M23_08890), however, was not in direct genetic vicinity of the RNF-gene cluster. Furthermore, we identified a second gene with homologies to *rseC* in *C. carboxidovorans* (Cca_07835) and *C. kluyveri* (CKL_2767), but neither RNF complex genes nor other genes that are involved in the autotrophic metabolism, such as the genes for the Wood-Ljungdahl pathway, are located in the direct vicinity of this second *rseC* homolog (Table 3). Notably, we also identified a *rseC* gene in the non-acetogenic bacterium *R. capsulatus*, which is the microbe in which the RNF complex was first described (12). The *rseC* gene in *R. capsulatus* is located upstream of *rnfF* instead of *rnfC*, which is separated by five genes (Fig. 5). Also *E. coli* possesses one *rseC* gene that is organized in the *rseABC* gene cluster (Fig. 5, **Supplementary Text S1D**) (19).

**Fig. 5.**
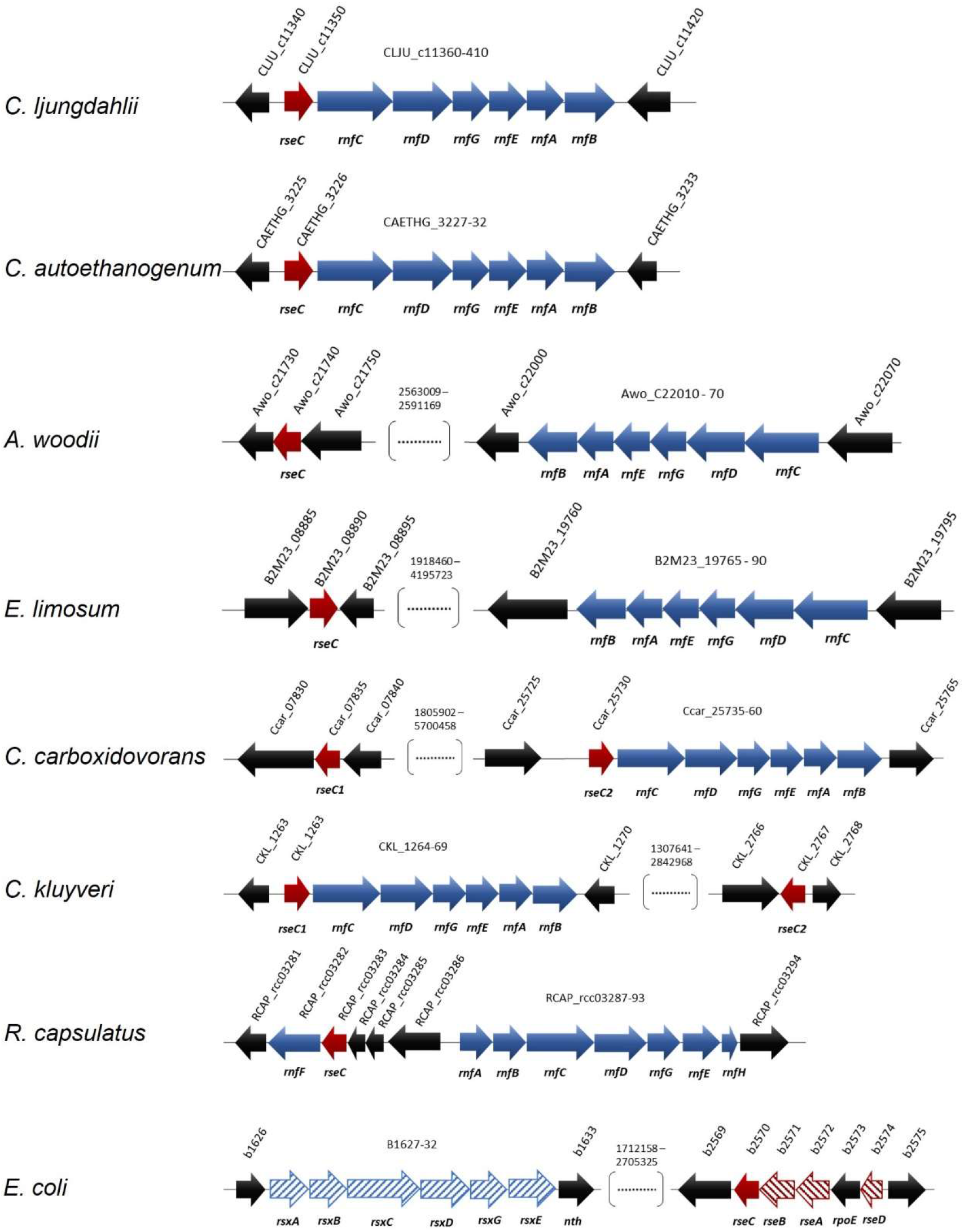
Location and orientation of *rseC* genes in microbes that possess RNF complex gene clusters. The conserved protein domain RseC_MucC (pfam04246) was identified in the *rseC* protein sequence of *C. ljungdahlii* and used to search for putative *rseC* genes in the genome of *C. autoethanogenum, A. woodii*, *E. limosum, C. carboxidovorans, C. kluyveri, R. capsulatus*, and *E. coli*. All sequence analyses and gene arrangements were adapted from the JGI platform and the NCBI database (03/2021). The type strains are listed in Table 3. In red, putative *rseC* genes; in red pattern fill, *rseC*-associated genes in *E. coli*; in blue, RNF-complex gene cluster; in blue pattern fill, *rsx* genes, which are homologous to the *rnf* genes in *R. capsulatus*.

**Table 3.** Distribution of *rseC* genes.

| microbe<sup>a</sup> | amount of<br>*rseC* genes<sup>b</sup> | RNF or Ech | *rseC* associated<br>with RNF genes | gene locus |
| --- | --- | --- | --- | --- |
| *Clostridium ljungdahlii* | 1 | RNF<sup>c</sup> | yes | CJLU_c11350 |
| *Clostridium autoethanogenum* | 1 | RNF<sup>c</sup> | yes | CAETHG_3226 |
| *Clostridium carboxidovorans* | 2 | RNF<sup>c</sup> | yes, one of them | Ccar_07835, Ccar_025730 |
| *Clostridium kluyveri* | 2 | RNF<sup>c</sup> | yes, one of them | CKL_1263, CKL_2767 |
| *Eubacterium limosum* | 1 | RNF<sup>d</sup> | no | B2M23_08890 |
| *Acetobacterium woodii* | 1 | RNF<sup>d</sup> | no | Awo_c21740 |
| *Thermotoga maritima* | 1 | RNF<sup>d</sup> | no | THEMA_1487 |
| *Moorella thermoacetica* | 0 | Ech | no | - |
| *Thermoanaerobacter kivui* | 0 | Ech | no | - |
| *Rhodobacter capsulatus* | 1 | RNF<sup>e</sup> | yes | RCAP_rcc03283 |
| *Escherichia coli* | 1 | Rsx<sup>f</sup> | no, but with Rsx | b2570 |

<sup>a</sup> The type strains were: *C. ljungdahlii* DSM13528; *C. autoethanogenum* DSM10061; *C. carboxidovorans* P7; *C. kluyveri* DSM555; *E. limosum* ATCC8486; *A. woodii* DSM1030; *T. maritima* DSM3109; *M. thermoacetica* ATCC39073; *T. kivui* DSM2030; *R. capsulatus* SB1003; and *E. coli* K-12.
<sup>b</sup> The pfam domain pfam04426 was used to search for putative *rseC* genes in each genome.
<sup>c</sup> The RNF complex uses (or is supposed to use) protons.
<sup>d</sup> The RNF complex uses (or is supposed to use) sodium ions.
<sup>e</sup> The RNF complex either uses protons or sodium ions. Experimental data are missing.
<sup>f</sup> Rsx is encoded by *rsxABCDGE* and is homologous to the RNF-gene cluster in *R. capsulatus*.

The conservation of the RseC amino-acid sequence was between 59% and 100% for *C. ljungdahlii*, *C. autoethanogenum*, *C. carboxidovorans*, and *C. kluyveri*, which is a high similarity (**Supplementary Table S2**, **Supplementary Fig. S5**). In addition, the amino-acid sequence length is nearly identical with 138 amino acids (*C. ljungdahlii*, *C. autoethanogenum*, and *C. carboxidovorans*) and 137 amino acids (*C. kluyveri*), respectively. The second RseC homolog from *C. carboxidovorans* and *C. kluyveri* shared an identity of 65% with each other, but only between 25% and 49% to all other RseC proteins (**Supplementary Table S2**, **Supplementary Fig. S5**). The RseC from *A. woodii* and *E. limosum* shared a similarity of 57% with each other, and only of 34% to 35% with the RseC proteins that are encoded directly upstream of the RNF-gene clusters (Fig. 5, **Supplementary Table S2**, **Supplementary Fig. S5**). The RseC proteins from *R. capsulatus* and *E. coli* have the same amino-acid sequence length (159 amino acids), but shared low similarities to each other (31%) as well as to the RseC proteins from the other microbes (18-34%) (**Supplementary Table S2**, **Supplementary Fig. S5**). The similarity of the RseC protein from *C. ljungdahlii* and *R. capsulatus* was only 23%, while it was 36% for the RseC protein from *C. ljungdahlii* in comparison to the RseC protein from *E. coli*. Overall, the RseC protein sequence seems to be highly conserved in acetogens that contain an RNF-gene cluster.

### The *nar* gene cluster encodes a functional nitrate reductase in *C. ljungdahlii*

We had found that nitrate reduction during heterotrophy is impacted for the *C. ljungdahlii* ΔRNF strain but not the *C. ljungdahlii* Δ*rseC* strain. Thus, we aimed to explore nitrate metabolism and the interplay with the RNF complex further. For *C. ljungdahlii*, it was postulated that nitrate is reduced by nitrate reductase to nitrite and, subsequently, converted *via* nitrite reductase and hydroxylamine reductase into ammonium, and the involved genes were predicted in the genome (14, 27). Emerson *et al.* (23) had found that in the presence of nitrate the expression level of the genes that encode the putative nitrate reductase (CLJU_c23710-30) were significantly increased. The three genes are annotated as nitrate reductase NADH oxidase subunit (CLJU_c23710), nitrate reductase electron transfer subunit (CLJU_c23720), and nitrate reductase catalytic subunit (CLJU_c23730) (14). We refer to these three genes (CLJU_c23710-30) as the *nar* gene cluster. We verified the absence of the *nar* gene cluster from the genome of the *C. ljungdahlii* Δ*nar* strain, after mediating the deletion with our CRISPR-Cas12a system (**Supplementary Fig. S2B**). This strain was able to grow during autotrophy and heterotrophy, but had completely lost the ability to reduce nitrate under both conditions (Fig. 6F, **Supplementary Fig. S6F**). We observed similar growth and pH behavior for cultures of *C. ljungdahlii* Δ*nar* during autotrophy with either ammonium or nitrate (Fig. 6A, 6B, **Supplementary Fig. S6A, S6B**). Enhanced autotrophic growth in nitrate-containing medium when compared to ammonium-containing medium, such as with the wild-type strain, was not detected (Fig. 6A).

**Fig. 6.**
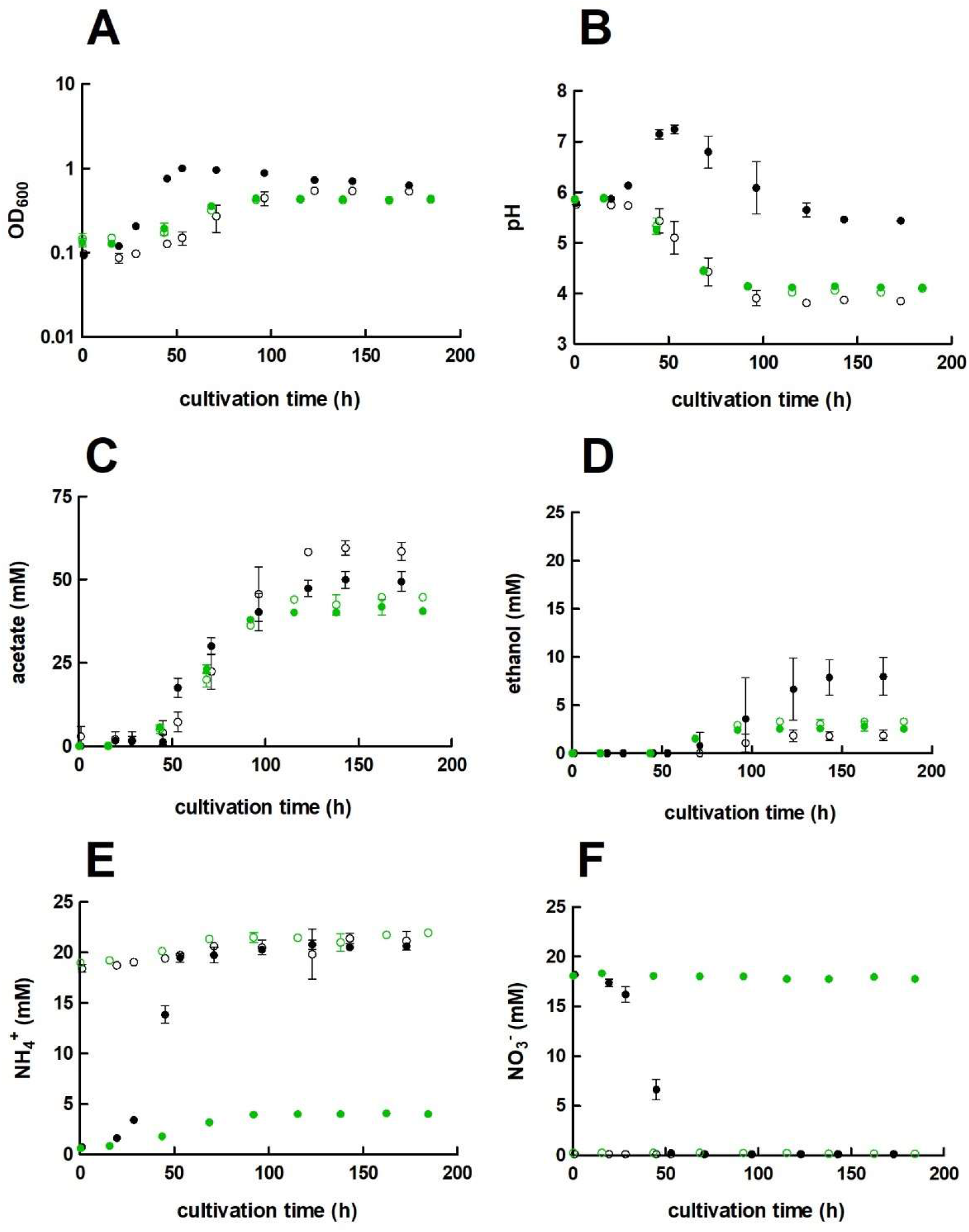
Growth, pH behavior, nitrate reduction of *C. ljungdahlii* Δ*nar* with H_2_ and CO_2_. Cultures were grown in 100 mL PETC medium in 1 L bottles at 37°C and 150 rpm for 185 h. The headspace consisted of H_2_ and CO_2_ (80/20 vol-%) and was set to 0.5 bar overpressure. The medium contained either 18.7 mM nitrate (NO_3_^-^) (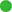) or 18.7 mM ammonium (NH_4_ ^+^) (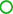) as nitrogen source. The *C. ljungdahlii* WT data (●, ○) from Supplementary Fig. S1 is given for comparison. All cultures were grown in biological triplicates, data is given as mean values, with error bars indicating the standard deviation. **A**, growth; **B**, pH-behavior; **C**, acetate concentrations; **D**, ethanol concentration; **E**, ammonium concentration; and **F**, nitrate concentrations. Δ*nar*, deletion of nitrate reductase gene cluster; rpm, revolutions per minute; CO_2_, carbon dioxide; and H_2_, hydrogen.

However, we still observed differences in the growth when compared to *C. ljungdahlii* WT. Growth rates during autotrophy of *C. ljungdahlii* Δ*nar* were 0.018 h^-1^ for ammonium- and 0.017 h^-1^ for nitrate-containing medium, which is a significant reduction of 24% (*P* = 0.04) and 76% (*P* ≤ 0.001) in comparison to the wild type (Table 1). The maximum observed OD_600_ values were both 0.44±0.01, which is a significant decrease of 21% (*P* ≤ 0.001) for ammonium cultures and 55% (*P* = 0.002) for nitrate cultures when compared to the wild type (Fig. 6A). A pH increase as a consequence of ammonium production, such as observed for the wild type, was not observed in cultures of *C. ljungdahlii* Δ*nar* (Fig. 6B). The maximum acetate concentrations were significantly reduced by 25% (44.8±0.2 mM, *P* ≤ 0.001) for ammonium cultures and by 16% (41.9±1.9 mM, *P* = 0.01) for nitrate cultures of *C. ljungdahlii* Δ*nar* (Fig. 6C, Table 1). The maximum ethanol concentrations were significantly increased by 79% (3.3±0.2 mM, *P* = 0.02) and significantly decreased by 64% (2.9±0.4 mM, *P* = 0.02) for ammonium and for nitrate conditions with carbon dioxide and hydrogen, respectively (Fig. 6D, Table 1). Even though *C. ljungdahlii* Δ*nar* was not able to use nitrate, cultures still accumulated 3-4 mM ammonium until the end of the cultivation in nitrate-containing medium (Fig. 6E).

The growth rates for heterotrophic cultures were 0.071 h^-1^ for ammonium- and 0.067 h^-1^ for nitrate containing medium (**Supplementary Table S1**). The maximum observed OD_600_ value was 2.35±0.04 for ammonium cultures of *C. ljungdahlii* Δ*nar*, which is similar to the performance of *C. ljungdahlii* WT (**Supplementary Fig. S6A**). For nitrate cultures the maximum observed OD_600_ value was 1.51±0.03 and corresponds to a significant reduction of 32% (*P* ≤ 0.001) when compared to *C. ljungdahlii* WT under the same conditions (**Supplementary Fig. S6A**). The maximum acetate concentrations were 51.9±0.9 mM for ammonium- and 28.7±1.1 for nitrate-containing medium, which is a reduction of 1% (*P* = 0.6) and a significant reduction of 34% (*P* ≤ 0.001), respectively (**Supplementary Fig. S6C**). Interestingly, the maximum ethanol concentrations for *C. ljungdahlii* Δ*nar* significantly increased by 45% (15.3±0.1 mM, *P* ≤ 0.001) when ammonium and fructose were provided, and by 234% (16.6±0.2, *P* ≤ 0.001) when nitrate and fructose were provided (**Supplementary Table S1**, **Supplementary Fig. S6D**). The provided fructose was only consumed completely by *C. ljungdahlii* Δ*nar* in ammonium-containing but not in nitrate-containing medium (**Supplementary Fig. S6G**).

Finally, we confirmed that the complementation of *C. ljungdahlii* Δ*nar* with the plasmid pMTL83152_*nar*, which encodes the *nar* gene cluster under the expression control of the constitutive P*_thl_* promoter, enabled the *C. ljungdahlii* Δ*nar* pMTL83152_*nar* strain to utilize nitrate under autotrophic conditions again, while this was not possible in an empty plasmid control strain (**Supplementary Fig. S7**). The nitrate cultures of *C. ljungdahlii* Δ*nar* pMTL83152_*nar* reached a growth rate of 0.054 h^-1^ and maximum observed OD_600_ values of 1.54±0.03, which is a significant reduction of 26% (*P* = 0.004) and a significant increase of 54% (*P* ≤ 0.001) in comparison to the wild type when growing with nitrate (Table 2). Maximum acetate concentrations were 41.7±2.5 mM, while maximum ethanol concentrations were 3.4±0.5 mM (**Supplementary Fig. S7C, S7D**). This is a significant reduction of 17% (*P* = 0.02) and of 57% (*P* ≤ 0.001) in contrast to the nitrate-grown cultures of *C. ljungdahlii* WT (Table 2). Therefore, we revealed that the expression of the *nar* gene cluster led to the only functional nitrate reductase in *C. ljungdahlii* under the tested conditions.

## Discussion

### A functional RNF complex is essential for autotrophy but not for heterotrophy in *C. ljungdahlii*

Here, we provided further insight into the autotrophy of *C. ljungdahlii* and the connection to nitrate metabolism. With the strain *C. ljungdahlii* ΔRNF, we confirmed that the absence of the RNF complex leads to a complete loss of autotrophy in *C. ljungdahlii*. Unlike in a previous study by Tremblay *et al.* (13), this strain provides a stable genotype that cannot revert back to the wild-type genotype, which can be used to further study the energy conservation principles in this acetogen (Fig. 1B, 2). Heterotrophic growth in this strain was still possible, but considerably reduced when compared to the wild type (Fig. 1C, **Supplementary Fig. S1**). While we did not measure the difference in the headspace gas composition during heterotrophy for *C. ljungdahlii* ΔRNF and wild type, we argue that *C. ljungdahlii* ΔRNF lost the ability to fixate the carbon dioxide that is released during glycolysis, which is the defining feature of acetogens (1, 10). Thus, even though the Wood-Ljungdahl pathway was still present, this strain was not able to balance the electrons in the metabolism to drive the Wood-Ljungdahl pathway. Further research is required to confirm this hypothesis. The RNF deletion in *A. woodii* did also lead to reduced acetate production during heterotrophy, but the strain reached similar OD_600_ values compared to the *A. woodii* wild type (25). In comparison to *C. ljungdahlii*, the RNF complex of *A. woodii* uses sodium ions instead of protons to generate the chemiosmotic gradient, which is then consumed by a sodium-dependent F_1_F_O_ ATPase to generate ATP (28, 29). Overall, this further confirms the meticulous differences in the energy conservation and redox balancing in different acetogens (2), which have to be considered to apply acetogens for biotechnological purposes.

### RseC is a positive regulator of the RNF complex genes and plays a critical role during autotrophy

We further investigated the regulation of the RNF-gene cluster by the putative regulator RseC. The *rseC* gene is known to encode a transcriptional regulator in other microbes such as *E. coli* and *S. typhimurium* (**Supplementary Text S1D**) (17, 19, 20, 30). Our results demonstrated that RseC played a critical role for the formation of a functional RNF complex in *C. ljungdahlii* (Table 2, Fig. 2). A deletion of the *rseC* gene led to the complete loss of autotrophy (Fig. 2). With our qPCR analyses, we confirmed that RseC, indeed, had a positive regulatory effect on the expression of the RNF-gene cluster during autotrophy. Our results indicate that RseC is essential for the activation of RNF-gene cluster expression during autotrophy, but not during heterotrophy, while we cannot rule out other modulating activities (Fig. 4A, 4C, 7, **Supplementary Fig. S4**, **Supplementary Text S1E**). Further biochemical and molecular biological investigations, such as the purification of the RseC protein and DNA-binding assays, or the study of the subcellular localization, will be required to unravel the regulatory functions of RseC in *C. ljungdahlii* and other acetogens with an RNF complex in more detail.

**Fig. 7.**
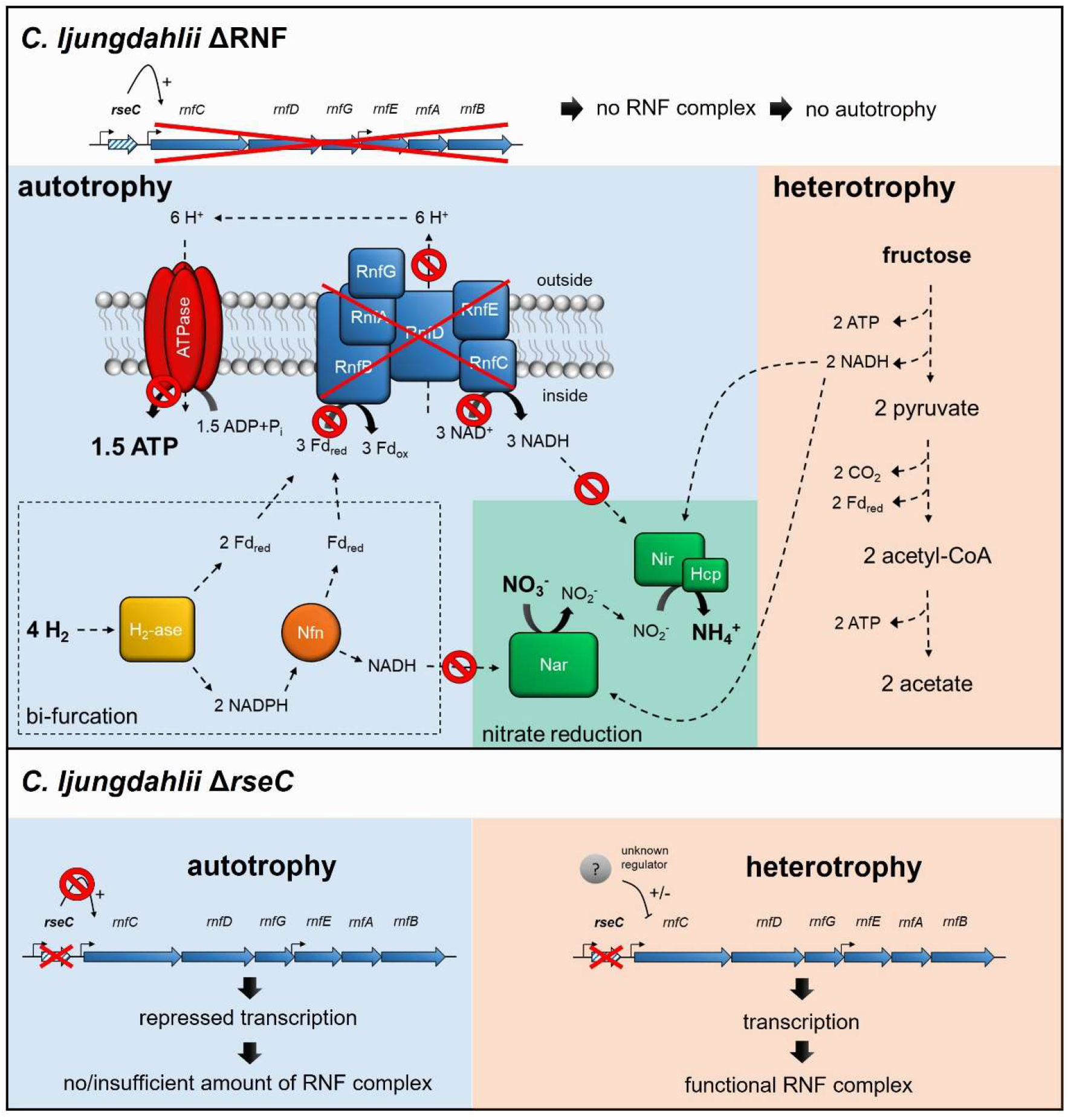
Schematic model of RNF-gene regulation and nitrate reduction in the deletion strains *C. ljungdahlii* ΔRNF and *C. ljungdahlii* Δ*rseC* during autotrophy and heterotrophy. In both deletion strains, nitrate reduction is not possible in non-growing cells during autotrophy with carbon dioxide and hydrogen due to the lack of a functional RNF complex, and thus the missing regeneration of reducing equivalents such as NADH. On the contrary, nitrate reduction can proceed in *C. ljungdahlii* ΔRNF during heterotrophy with NADH, which is provided by glycolysis of fructose. In *C. ljungdahlii* Δ*rseC*, the RNF complex genes are repressed during autotrophy but not during heterotrophy, which indicates a further unknown regulation mechanism during heterotrophy. Thus, a functional RNF complex is formed, and nitrate reduction can proceed such as proposed for the wild type. Abbreviations: H_2_, hydrogen; H^+^, proton, CO_2_, carbon dioxide; NO_3_^-^, nitrate; NO_2_^-^, nitrite; NH_4_ ^+^, ammonium; ATP, adenosine triphosphate; ADP + P_i_, adenosine diphosphate + phosphate; Fd_red/ox_, reduced/oxidized ferredoxin; NADH/NAD^+^, reduced/oxidize nicotinamide adenine dinucleotide; NADPH/NADP^+^, reduced/oxidized nicotinamide adenine dinucleotide phosphate; RnfCDGEAB, RNF-complex subunits; Nar, nitrate reductase; Nir, nitrite reductase; Hcp, hydroxylamine reductase; H_2_-ase, bifurcating hydrogenase/lyase; Nfn, bifurcating transhydrogenase; e^-^, electron; ΔRNF, *C. ljungdahlii* ΔRNF; and Δ*rseC*, *C. ljungdahlii* Δ*rseC*. The model was adapted from Emerson *et al.* (23).

### Nitrate reduction does not require a functional RNF complex but benefits from a correct electron balance

Furthermore, we investigated the nitrate metabolism in *C. ljungdahlii*. We confirmed that the genes CLJU_c23710-30 encode the functional subunits of the only nitrate reductase under the tested conditions for *C. ljungdahlii* (Fig. 6, **Supplementary Fig. S6, S7**). In the presence of nitrate, *C. ljungdahlii* WT quickly utilized all nitrate even though a sufficient amount of nitrogen-source was covered by the added yeast extract (Fig. 2F, **Supplementary Fig. SF1**). Thus, nitrate reduction in *C. ljungdahlii* is mainly used for energy conversion, and therefore must be of a dissimilatory function (23) (**Supplementary Text S1F**). The stoichiometry for nitrate reduction in *C. ljungdahlii* is proposed as follows: 4 H_2_ + 2 H^+^ + NO_3_^-^+ 1.5 ADP + 1.5 P_i_ ⇌ 4 H_2_O + NH_4_^+^ + 1.5 ATP with *Δ*_r_G′_0_ = −150 kJ/mol H_2_ (23, 31). This mechanism would require electron bifurcation from the hydrogenases and the activity of the RNF complex, but would then provide ATP completely independent of the Wood-Ljungdahl pathway (or more general, independent of the carbon metabolism) (23, 32). Thus, we hypothesized that nitrate reduction in *C. ljungdahlii* requires a functional RNF complex for a correct electron balance. Indeed, non-growing cells of both *C. ljungdahlii* ΔRNF and *C. ljungdahlii* Δ*rseC* were not able to reduce nitrate during autotrophy (Fig. 2F). However, nitrate reduction still proceeded in both deletion strains during heterotrophy (**Supplementary Fig. S1F**). In *C. ljungdahlii* ΔRNF a functional RNF complex was not present during heterotrophy because the RNF-complex encoding genes were deleted, but the required reducing equivalents for nitrate reduction were likely provided by glycolysis (Fig. 7). In contrast, in *C. ljungdahlii* Δ*rseC*, nitrate reduction was not impacted during heterotrophy, because the RNF complex genes were not repressed under these conditions and a functional RNF complex was formed (Fig. 4, 7). It remains to be answered whether there is a direct interplay between the nitrate reductase and the RNF complex, and whether this interplay is different during heterotrophy and autotrophy.

### The electron balance in the deletion strains is impacted beyond nitrate reduction

In general, the reduced growth indicated that *C. ljungdahlii* ΔRNF was not able to balance the electrons from glycolysis efficiently during heterotrophy. This led to the reduction in biomass and acetate production, while ethanol production was completely absent in heterotrophic cultures of *C. ljungdahlii* ΔRNF, which indicates that reducing power for a further reduction of acetate was not available (**Supplementary Table S1**, **Supplementary Fig. S1D**). In the batch experiments of Emerson *et al.* (23), *C. ljungdahlii* WT did not produce considerable amounts of ethanol when growing with nitrate (and carbon dioxide and hydrogen). When *C. ljungdahlii* WT was cultivated in pH-controlled bioreactors under continuous conditions, enhanced biomass and increased ethanol production rates were observed (24). This observation could not be fully explained yet, but it was assumed that electrons are predominantly used for the reduction of nitrate rather than for the reduction of acetate. This distribution of electrons changed in the absence of the nitrate reductase in the *C. ljungdahlii* Δ*nar* strain and higher maximum ethanol concentrations were observed (**Supplementary Text S1G**). It remains elusive, how the change in the distribution of electrons affects other NADH-dependent metabolic pathways in more detail. While further research is required to understand the regulatory mechanisms during autotrophy and the mechanism of energy conservation during nitrate reduction, with this work, we provide a deeper insight into the autotrophic metabolism and nitrate reduction in *C. ljungdahlii*.

## Methods

### Bacterial strains and growth

*Escherichia coli* TOP10 (Invitrogen), *E. coli* EPI300 (Lucigen), and *E. coli* HB101 PKR2013 (DSM 5599) were grown at 37°C in Luria Broth (LB) medium containing (per liter): 5 g NaCl; 10 g peptone; and 5 g yeast extract. *C. ljungdahlii* ATCC13528 was generally cultivated in anaerobic Rich Clostridial Medium (RCM) containing per liter: 5 g fructose; 3 g yeast extract; 10 g meat extract; 10 g peptone; 5 g NaCl; 1 g soluble starch; 3 g sodium acetate; 0.5 g L-cysteine HCl; and 4 mL resazurin-solution (0.025 vol-%). For growth experiments with *C. ljungdahlii*, standard PETC medium (24) was used containing (per liter): 1 g yeast extract; 1.0 g NH_4_Cl; 0.1 g KCl; 0.2 g MgSO_4_x7 H_2_O; 0.8 g NaCl; 0.1 g KH_2_PO_4_; 0.02 g CaCl_2_x2 H_2_O; 4 mL resazurin-solution (0.025 vol-%); 10 mL trace element solution (TE, 100x); 10 mL Wolfe’s vitamin solution (100x); 10 mL reducing agent (100x); and 20 mL of fructose/2-(N-morpholino)ethanesulfonic acid (MES) solution (50x). TE was prepared as 100x stock solution containing (per liter): 2 g nitrilotriacetic acid (NTA); 1 g MnSO_4_xH_2_O; 0.8 g Fe(SO_4_)2(NH_4_Cl)2×6 H_2_O; 0.2 g CoCl_2_x6 H2O; 0.0002 g ZnSO_4_x7 H_2_O; 0.2 g CuCl_2_x2 H_2_O; 0.02 g NiCl_2_x6 H_2_O; 0.02 g Na_2_MoO_4_x2 H_2_O; 0.02 g Na_2_SeO_4_; and 0.02 g Na_2_WO_4_. The pH of the TE was adjusted to 6.0 after adding NTA. The solution was autoclaved and stored at 4°C. Wolfe’s vitamin solution was prepared aerobically containing (per liter): 2 mg biotin; 2 mg folic acid; 10 mg pyridoxine-hydrochloride; 5 mg thiamin-HCl; 5 mg riboflavin; 5 mg nicotinic acid; 5 mg calcium pantothenate; 5 mg p-aminobenzoic acid; 5 mg lipoic acid; and 0.1 mg cobalamin. The vitamin solution was sterilized using a sterile filter (0.2 µm), sparged with N_2_ through a sterile filter, and stored at 4°C. The 50x fructose/MES solution contained (per 100 mL): 25 g fructose; and 10 g MES. The pH was adjusted to 6.0 by adding KOH. For autotrophic experiments, fructose was omitted. In nitrate experiments, ammonium chloride was replaced with sodium nitrate (NaNO_3_) in the equal molar amount (=18.7 mM). The reducing agent solution contained (per 100 mL): 0.9 g NaCl and 4 g L-cysteine HCl and was prepared with anaerobic water under anaerobic conditions. The reducing agent was stored at room temperature. For solid LB medium, 1.5 weight-% agar was added. For solid RCM or PETC medium 1.0-2.0 weight-% agar was added. For conjugation of *C. ljungdahlii* cells (see below) a modified PETC medium (PETC+5gS) was used containing additionally (per liter): 5 g peptone and 5 g meat extract.

Liquid *E. coli* cultures and autotrophic *C. ljungdahlii* cultures were agitated at 150 revolutions per minute (rpm) (Lab Companion Incubater Shaker ISS-7100R, Jeio Tech). Heterotrophic cultures of *C. ljungdahlii* and LB plates with *E. coli* cells were incubated without shaking (Incubator IN260, Memmert). Anaerobic work was performed in an anaerobic chamber (Glovebox-System UNIlab Pro, MBraun) with an N_2_ (100 vol-%) atmosphere. However, *C. ljungdahlii* cultures in bottles were transferred at the bench with sterile syringes and needles. Before each transfer between serum bottles, we flamed the top of the rubber stopper with ethanol (70 vol-%) at a Bunsen burner. All plating work with *C. ljungdahlii* was performed in the anaerobic chamber with a maximum of 5 parts per million (ppm) oxygen in the atmosphere. All plating work with *E. coli* was carried out in a lamina flow bench (Hera Safe KS18, Thermo Fischer Scientific). Antibiotics (see below) were added to maintain plasmid stability in recombinant cultures of *E. coli* and *C. ljungdahlii*.

### Antibiotics

Chloramphenicol (30 mg/mL), ampicillin (100 mg/mL), and kanamycin (50 mg/mL) were applied to maintain plasmids in *E. coli* strains, while thiamphenicol (5 mg/mL) was used for recombinant strains of *C. ljungdahlii*. Thiamphenicol was prepared as aerobic stock solution (25 mg/mL) in DMSO (100 vol-%) and diluted with sterile water (1:10) before use. The diluted thiamphenicol solution (2.5 mg/mL) was transferred into a sterile 1 mL syringe. 100 µL of this solution was used to add to a 50 mL RCM or PETC medium (final concentration of 5 mg/mL). The use of DMSO over ethanol as solvent for thiamphenicol prevented the addition of external ethanol to cultures of *C. ljungdahlii*, which is a metabolite. The thiamphenicol stock solution was stored at −20°C.

### Design and generation of CRISPR-FnCas12a plasmids for gene deletion

The broad-host plasmid pMTL83152 (Heap et al. 2009) was used as backbone (Table 4). All PCR steps were performed with Q5® High-Fidelity DNA Polymerase (New England Biolabs) and primers provided by IDT (Integrated DNA Technologies) (Table 5). PCR products were purified with QIAquick PCR Purification Kit (Qiagen). The gene *Fncas12a* of *Francisella novicida* (Zetche et al. 2015) was obtained from plasmid pY001 (Addgene #69973) and amplified with primers cas12a_fwd_BamHI and cas12a_rv_NcoI generating *Bam*HI and *Nco*I restriction sites for a subsequent restriction cloning to generate pMTL83152::*Fn*Cas12a. Two homology-directed repair arms (HDR1/HDR2) each with a size of 1000-1200 bp, which flank the targeted gene, were individually amplified with HDR_upst_fwd/rv and HDR_dwst_fwd/rv primers generating an overlap of 25-40 bp to each other. The fragments were purified, and 50-100 ng of both fragments were used as template for a subsequent fusion PCR using HDR_upst_fwdOv and HDR_dwst_rvOv primers, which generated new overlaps at 5’ and 3’ (fusion fragment HDR1/2). An crRNA array was synthesized and cloned as minigene into plasmid pUC19 by IDT (Integrated DNA Technologies) (Table 6). The crRNA array sequence contained the mini-promoter P4 (5′-TTGACAAATTTATTTTTTAAAGTTAAAATTAAGTTG-3′) (33), the FnCas12a-specific directed repeats (DR) sequence (5’-TAATTTCTACTGTTGTAGAT-3’) (34), 1-2 sgRNA for the targeted gene(s) (Pam sequence TTV for target RNF and TTTV for target *rseC* and *nar*), and the rrbn-T1-terminator (35). The crRNA array fragment was amplified with primers minigene_crRNA_fwd/rv creating overhangs to the fused HDR1/2 fragment and the plasmid backbone. For gene targets with a size >2 kb, such as *rnfCDGEAB* and *nar*, two sgRNA (and two DRs) were used in the same crRNA array (Table 6). For the assembly reaction (Gibson Assembly Ultra Kit, Synthetic Genomics), the plasmid pMTL83152::*Fncas12a* was first digested using *BbvC*I and CIP (New England Biolabs) for 3h at 37°C, purified by PCR-clean, and then mixed with the purified fused HDR1/2 fragment and the crRNA array fragment. Using electrocompetent E. coli EPI300 cells (TransforMax^TM^, Lucigen) and electroporation for transformation highly increased cloning efficiency for the CRISPR-Cas12a constructs in *E. coli*. For inducible Cas12a expression, the P*_thl_* module was replaced with the *tetR-O1* promoter module (P*_tetR-O1_*) (36) using restriction sites *Sbf*I and *Bam*HI, for all generated CRISPR-Cas12a plasmids.

**Table 4.** Plasmids used in this study.

| plasmid | function | source |
| --- | --- | --- |
| pMTL83151 | shuttle-vector | (37) |
| pMTL83152 | shuttle-vector with constitutive thiolase promoter $P_{thl}$ | (37) |
| pMTL2tetO1gusA | pMTL82254 with pminiThl:tetR and p2tetO1:gusA | (36) |
| pMTL8315tet | shuttle-vector with inducible promoter system *tetR-O1* | this study |
| pMTL83151_$P_{nat}$_rnfCDGEAB | overexpression of *rnfCDGEAB* through native promoter $P_{nat}$ | this study |
| pMTL83152_rseC | overexpression of *rseC* through constitutive promoter $P_{thl}$ | this study |
| pMTL83152_nar | overexpression of *nar* through constitutive promoter $P_{thl}$ | this study |
| pY001_FnCpf1(Cas12a) | expression of FnCas12a | (34), Addgene 69973 |
| pMTL83152_FnCas12a | constitutive expression of FnCas12a through $P_{thl}$ | this study |
| pMTL83152_FnCas12a_ΔrseC | constitutive expression of FnCas12a through $P_{thl}$, constitutive expression of a single sgRNA targeting *rseC* on the genome, fused repair HDR1/2 fragment for homologous recombination and marker-less gene deletion | this study |
| pMTL83152_FnCas12a_Δnar | constitutive expression of FnCas12a through $P_{thl}$, constitutive expression of two sgRNA targeting *nar* on the genome, fused repair HDR1/2 fragment for homologous recombination and marker-less gene deletion | this study |
| pMTL83152_FnCas12a_ΔrnfCDGEAB | constitutive expression of FnCas12a through $P_{thl}$, constitutive expression of two sgRNA targeting *rnfCDGEAB* on the genome, fused repair HDR1/2 fragment for homologous recombination and marker-less gene deletion | this study |
| pMTL8315tet_FnCas12a | inducible expression of FnCas12a through tetR-O1 promoter system | this study |
| pMTL8315tet_FnCas12a_ΔrseC | inducible expression of FnCas12a through tetR-O1 promoter system, constitutive expression of a single sgRNA targeting *rseC* on the genome, fused repair HDR1/2 fragment for homologous recombination and marker-less gene deletion | this study |
| pMTL8315tet_FnCas12a_Δnar | inducible expression of FnCas12a through tetR-O1 promoter system, constitutive expression of two sgRNA targeting *nar* on the genome, fused repair HDR1/2 fragment for homologous recombination and marker-less gene deletion | this study |
| pMTL8315te_FnCas12a_ΔrnfCDGEAB | inducible expression of FnCas12a through tetR-O1 promoter system, constitutive expression of two sgRNA targeting *rnfCDGEAB* on the genome, fused repair HDR1/2 fragment for homologous recombination and marker-less gene deletion | this study |

**Table 5.**
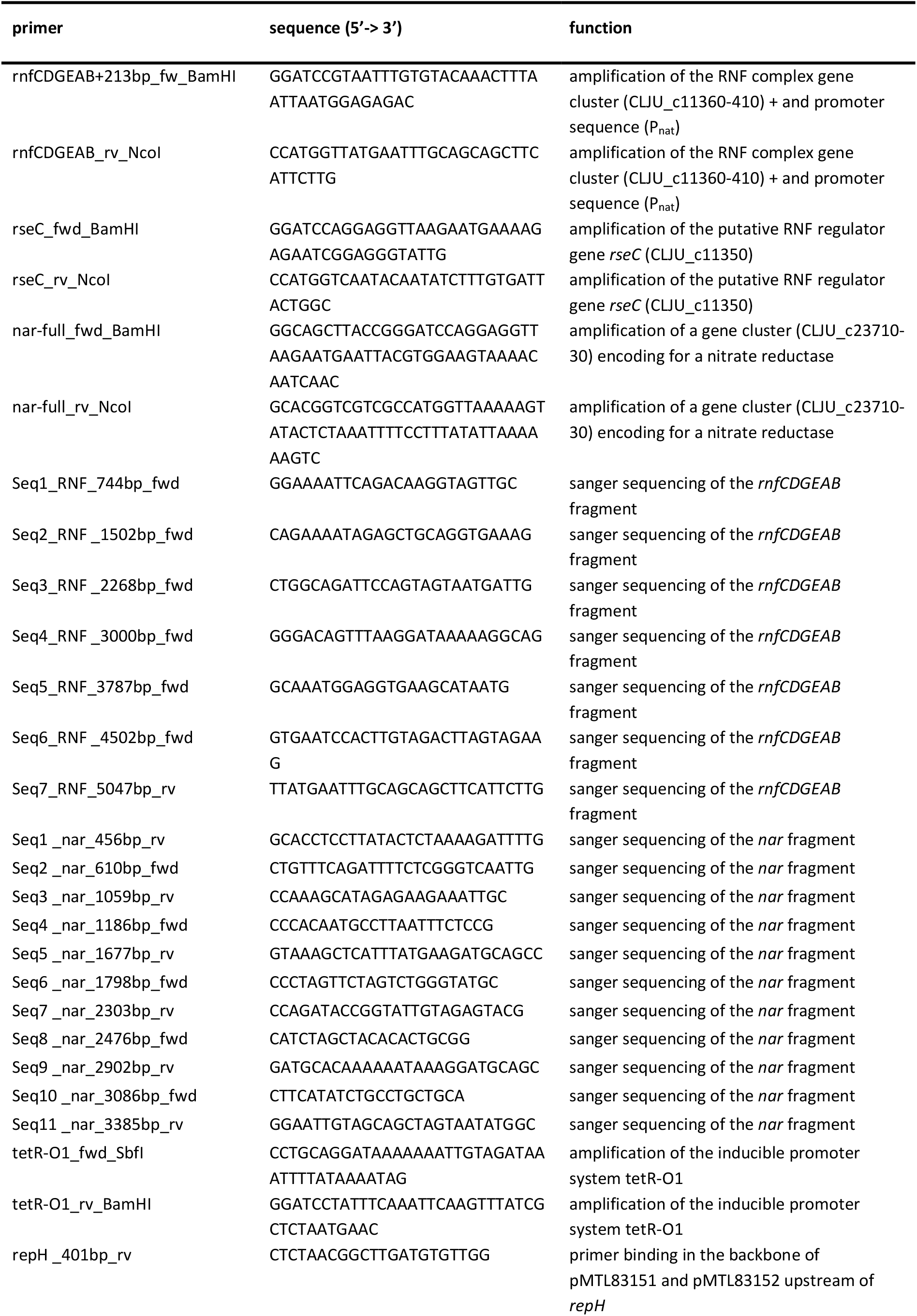
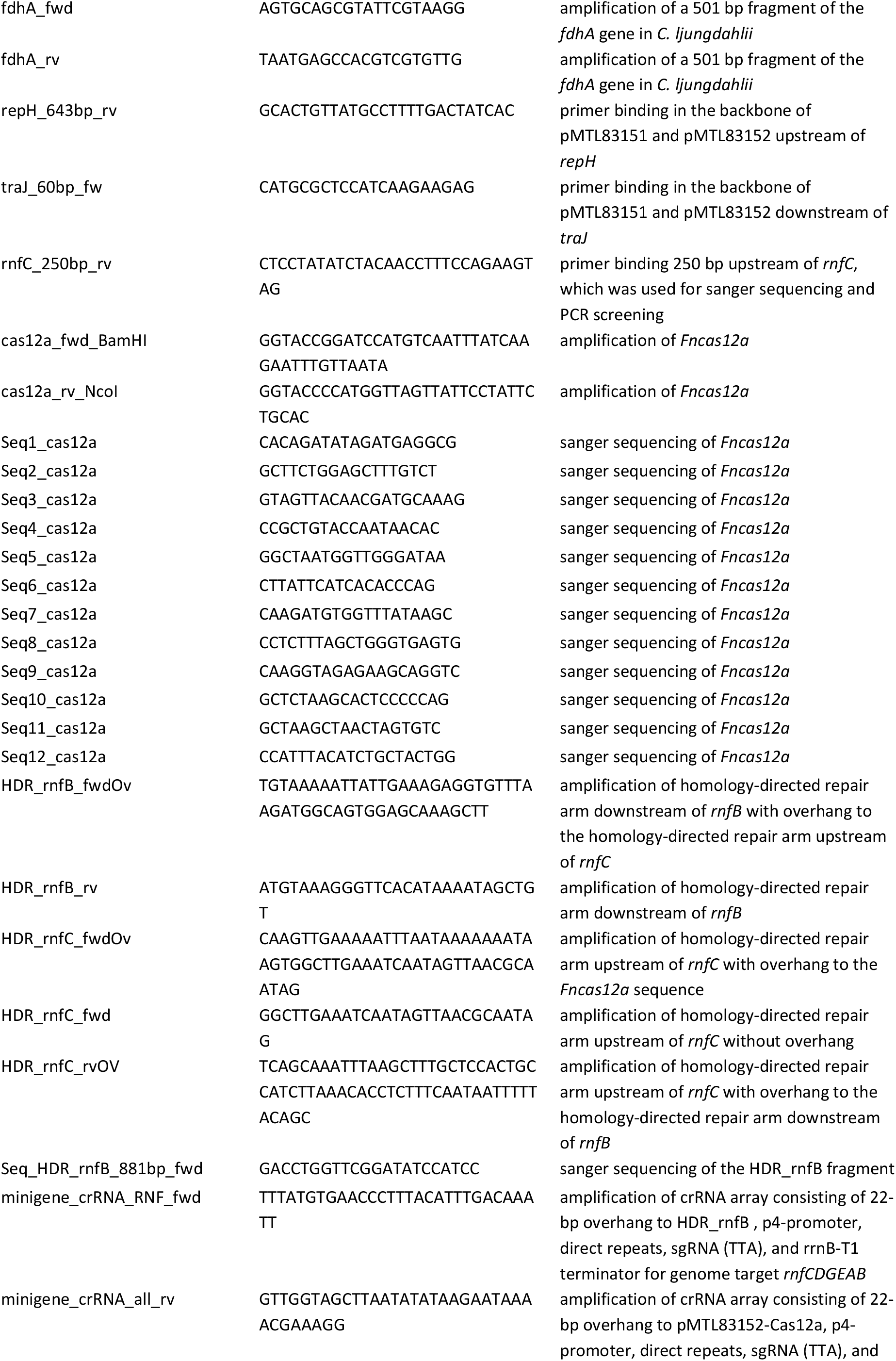
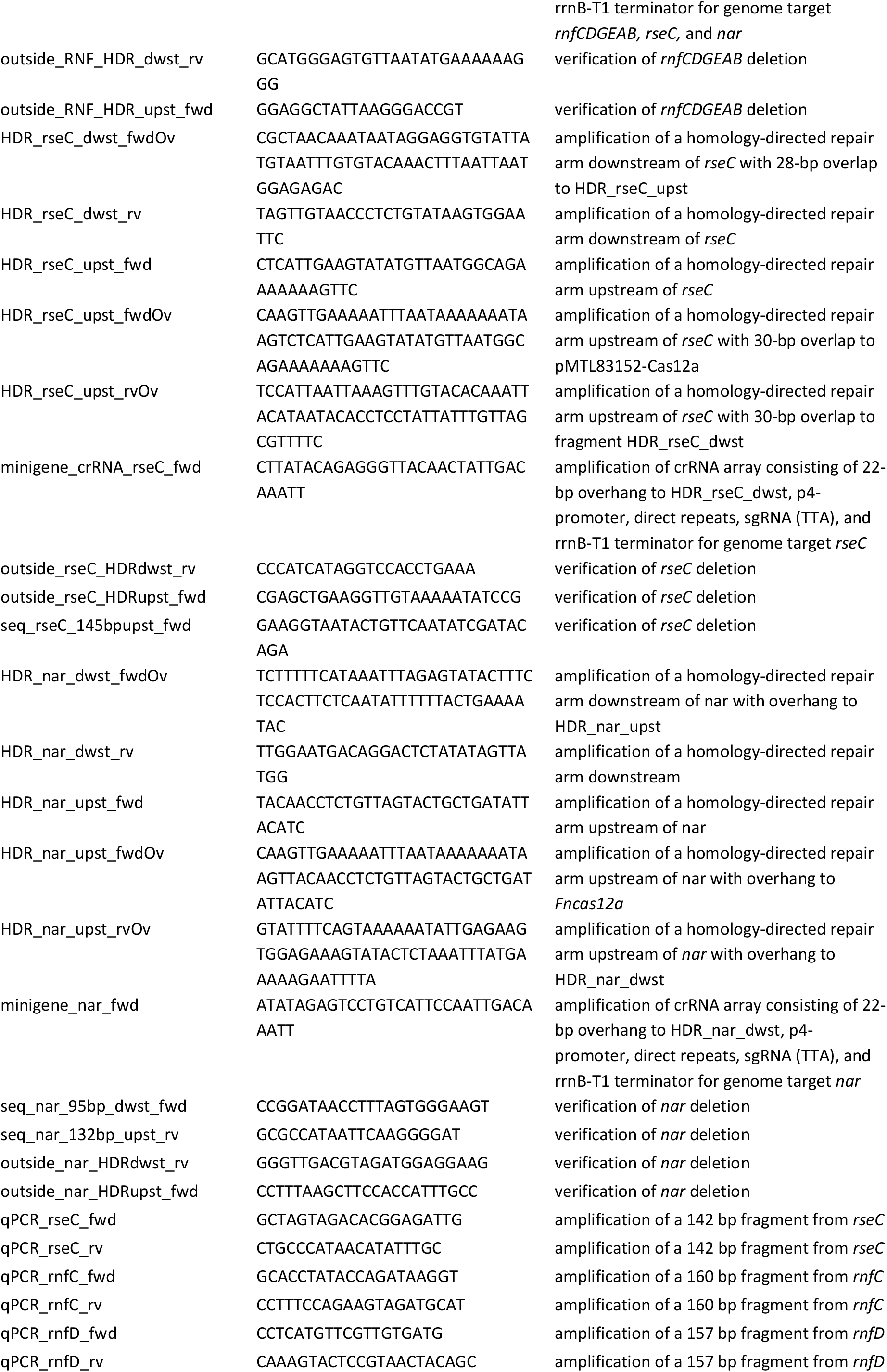
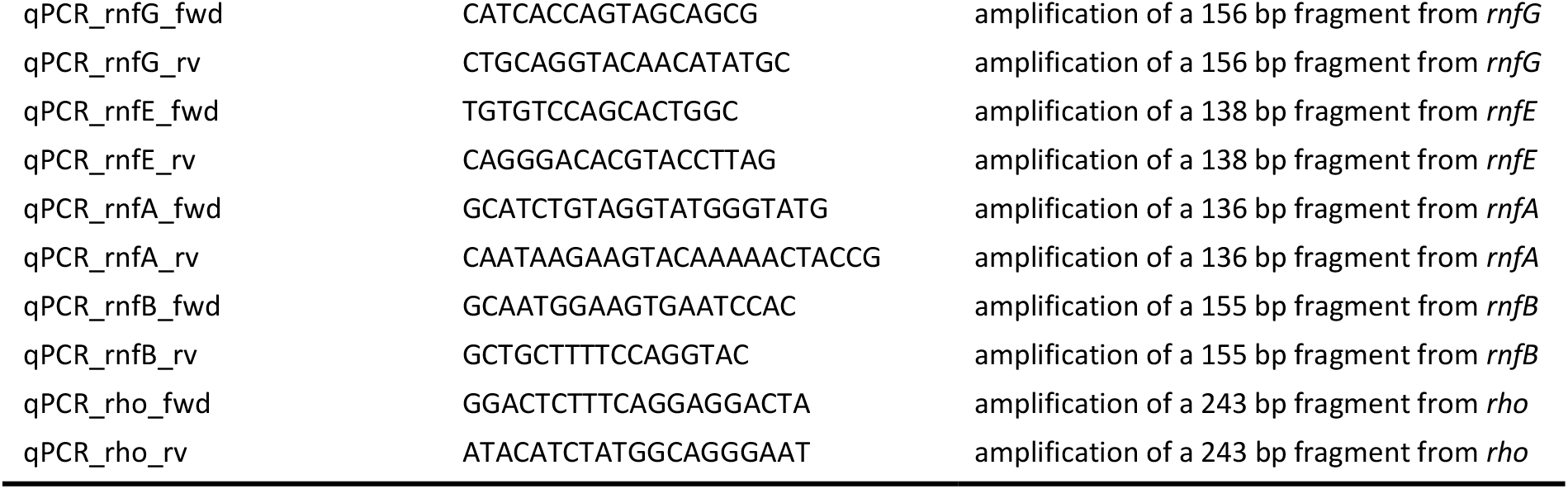
Primers used in this study.

**Table 6.**
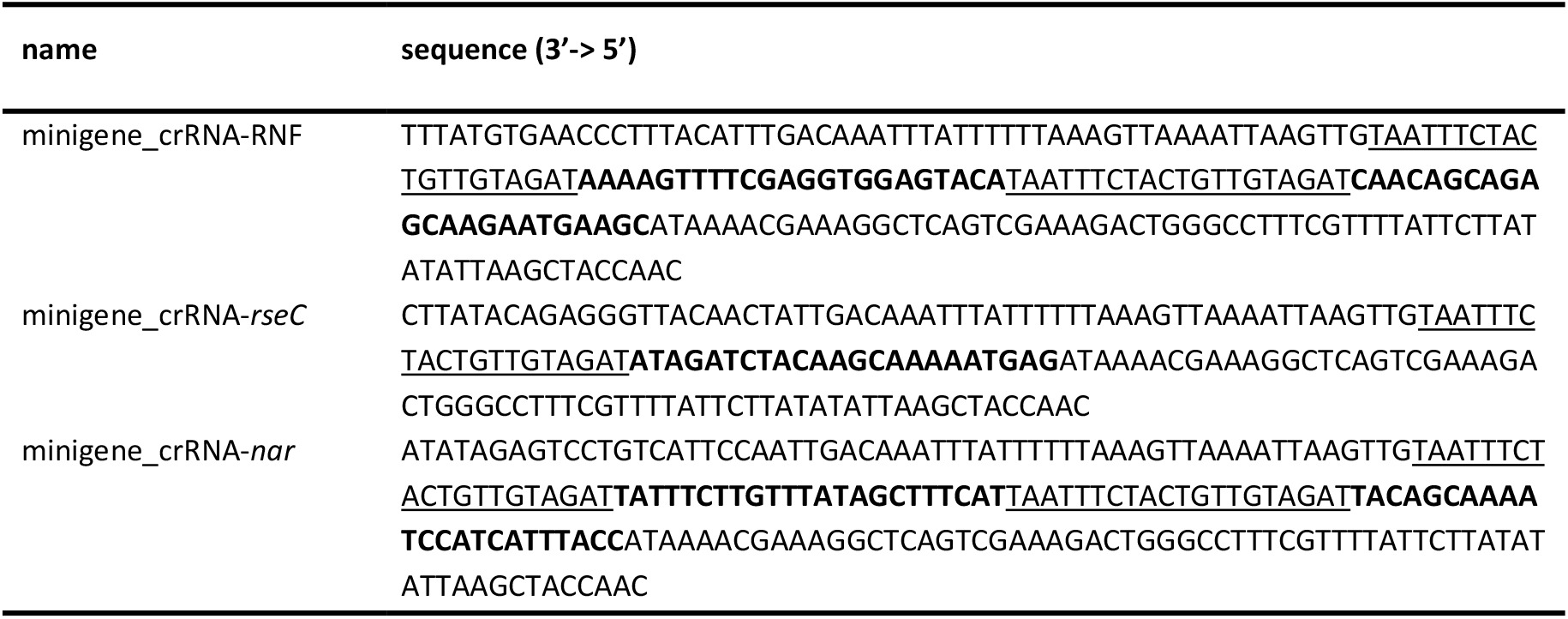
Synthesized mini genes that contain crRNA arrays for this study. Gene synthesis was performed by IDT (Integrated DNA Technologies). Each mini gene contains 20-22-bp overhang to the pMTL-backbone and to the fused homology-directed repair arms. Directed-repeats sequence of 20 bp (underlined). sgRNA with TTV PAM for the RNF complex gene cluster deletion and with TTTV PAM for the *nar* and *rseC* deletion (bold). Two sgRNAs were used to target RNF and *nar*.

### Generation of overexpression and complementation plasmids

The broad-host shuttle-vector system pMTL80000 (37) was used for all cloning steps. All generated plasmids of this study (Table 4) were cloned with restriction endonucleases and T4 ligase from NEB (New England Biolabs) or Gibson assembly (NEBuilder® HiFi DNA Assembly, New England Biolabs). PCR work was carried out with primers provided by IDT (Integrated DNA Technologies) (Table 5) and with a proof-reading Q5® High-Fidelity DNA Polymerase (New England Biolabs) according to the manufacture’s guidelines. Genomic DNA (gDNA) was purified from 2 mL of exponential cultures of *C. ljungdahlii* with the NucleoSpin Tissue Mini kit (Macherey-Nagel) and used as PCR-template. Notably, instead of performing harsh cell disruption according to the manufacture’s recommendation, we applied a 6×10 sec vortex interval during the procedure. The *rnfCDGEAB* gene cluster (CLJU_c11360-410) and a 213-bp sequence located upstream of *rnfC*, which contains the putative native promoter sequence (P_nat_), were amplified as one fragment using primers rnfCDGEAB+213bp_fwd and rnfCDGEAB_rv. The *rseC* gene (CLJU_c11350) was amplified using primers *rseC*_fwd and *rseC*_rv. The gene cluster CLJU_c23710-30, here referred to as *nar*, was amplified as one fragment using primers nar-full_fwd and nar-full_rv. All PCR products were purified with the QIAquick PCR Purification kit (Qiagen). Subsequently, the purified fragments were ligated into pMinit2.0 (New England Biolabs) and used for transformation of CaCl_2_-competent *E. coli* TOP10 cells (38). Next, the plasmid DNA was digested using the restriction sites determined by the used PCR primers and the fragment was cloned into the pMTL83151 plasmid generating pMTL83151_P_nat__*rnfCDGEAB* or into the pMTL83152 plasmid generating pMTL83152_*rseC* and pMTL83152_*nar*. Subsequently, all cloned fragments were verified again with test-digestion of the plasmid DNA and Sanger sequencing to exclude mutations in the gene sequences.

### Screening for correct plasmid DNA and genome editing

For screening and continuous purity control of our *C. ljungdahlii* strains (Table 7), we performed PCRs from culture samples or from purified DNA with the Phire Plant Direct PCR Master Mix (Thermo Fischer Scientific). *E. coli* colonies grown on selective LB plates after receiving plasmid constructs were analyzed for the correctly assembled plasmids using the Phire Plant Direct PCR Master Mix (Thermo Fischer Scientific). A small amount of recombinant *E. coli* cell material was directly transferred to the reaction mix. For *C. ljungdahlii* cells, 0.5-1 mL culture sample was harvested by centrifugation for 3 min at 13806 rpm (Centrifuge 5424, FA-45-24-11, Eppendorf) and resuspended in 100-500 µL 10 mM NaOH depending on the size of the cell pellet. Subsequently, cell suspensions were boiled for 10 min at 98°C. The hot reaction tubes were incubated on ice for 1 min and quickly vortexed before they served as a DNA template. In general, we used 20 µL PCR master mix, which consisted of 10 µL Phire Plant Mix 2x, 0.8 µL of each primer, 1 µL cell lysate sample, and 7.4 µl nuclease-free water. The PCR reaction was carried out according to the manufacture’s guidelines. We generally used the primers tra60bp_fwd and repH_401bp_rv or repH_643bp_rv for these control PCRs (Table 5), because they bind to the plasmid backbone of every pMTL plasmid used in this study. Verification of gene deletion in the genome of *C. ljungdahlii* was performed with “outside” primers, which bound upstream and downstream of the used homology-directed repair arms (HDR1/2) on the genomic DNA (Table 5). In addition, we performed test-digestion of the generated plasmids with restriction enzymes (New England Biolabs), and analyzed the fragment pattern *via* gel electrophoresis. The final plasmid sequence was verified by Sanger sequencing. Plasmid DNA was purified from *E. coli* with hand-made purification buffers (described below). Correct plasmid DNA was then purified with the QIAprep Spin Miniprep kit (Qiagen) *prior* to further use.

**Table 7.**
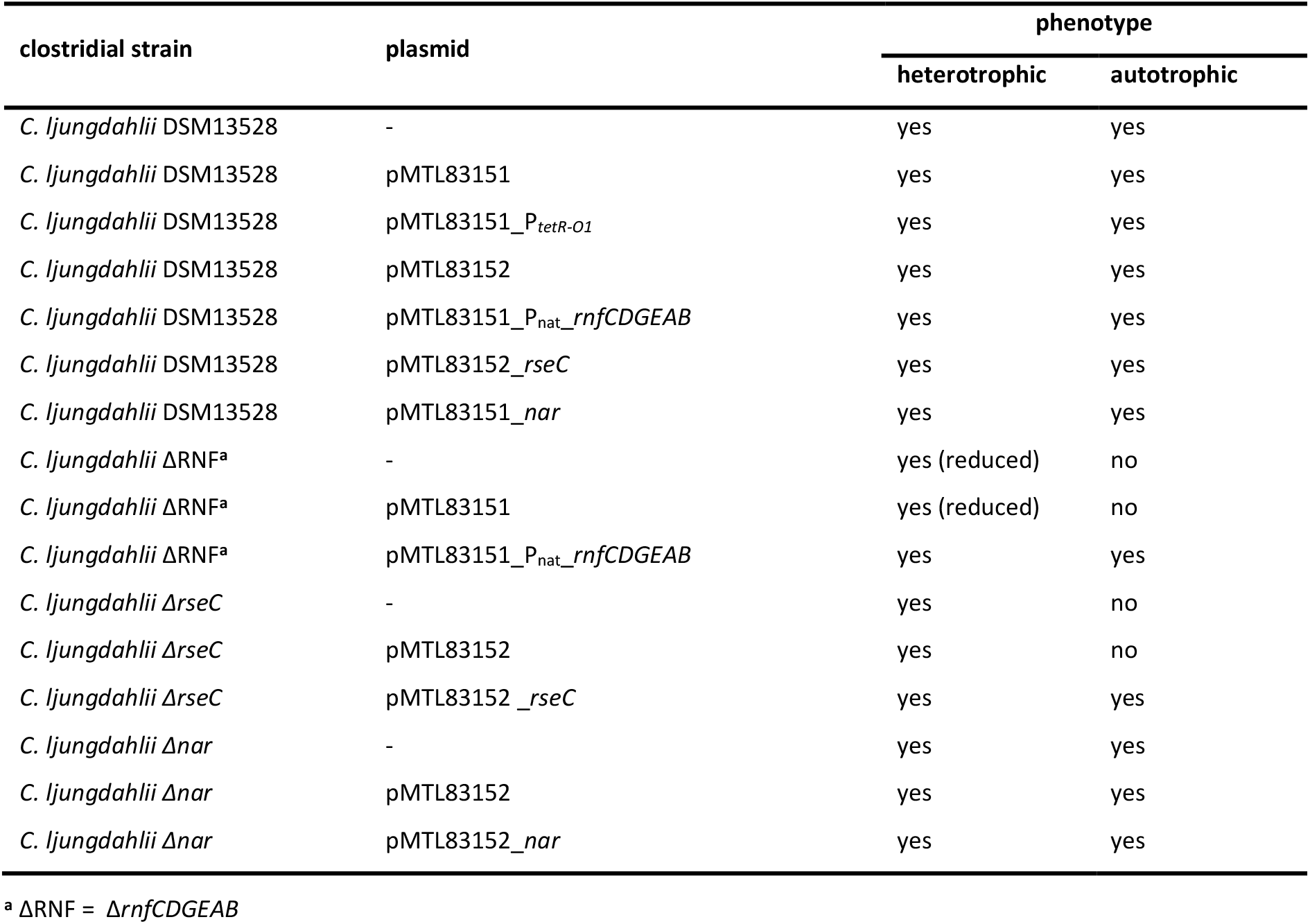
Used and generated C. ljungdahlii strains in this study.

| clostridial strain | plasmid | phenotype |  |
| --- | --- | --- | --- |
|  |  | heterotrophic | autotrophic |
| *C. ljungdahlii* DSM13528 | - | yes | yes |
| *C. ljungdahlii* DSM13528 | pMTL83151 | yes | yes |
| *C. ljungdahlii* DSM13528 | pMTL83151_P<sub>tetR-O1</sub> | yes | yes |
| *C. ljungdahlii* DSM13528 | pMTL83152 | yes | yes |
| *C. ljungdahlii* DSM13528 | pMTL83151_P<sub>nat_rnfCDGEAB</sub> | yes | yes |
| *C. ljungdahlii* DSM13528 | pMTL83152_rseC | yes | yes |
| *C. ljungdahlii* DSM13528 | pMTL83151_nar | yes | yes |
| *C. ljungdahlii* ΔRNF<sup>a</sup> | - | yes (reduced) | no |
| *C. ljungdahlii* ΔRNF<sup>a</sup> | pMTL83151 | yes (reduced) | no |
| *C. ljungdahlii* ΔRNF<sup>a</sup> | pMTL83151_P<sub>nat_rnfCDGEAB</sub> | yes | yes |
| *C. ljungdahlii* ΔrseC | - | yes | no |
| *C. ljungdahlii* ΔrseC | pMTL83152 | yes | no |
| *C. ljungdahlii* ΔrseC | pMTL83152_rseC | yes | yes |
| *C. ljungdahlii* Δnar | - | yes | yes |
| *C. ljungdahlii* Δnar | pMTL83152 | yes | yes |
| *C. ljungdahlii* Δnar | pMTL83152_nar | yes | yes |

<sup>a</sup> ΔRNF = ΔrnfCDGEAB

### A fast method for plasmid purification from *E. coli* without use of a commercial kit

For screening of successfully transformed *E. coli* cells we used a time- and money-saving protocol to purify plasmid DNA from multiple samples without using a commercial kit, which is a modified alkaline lysis protocol adapted from (39). All centrifugation steps were performed at 13806 rpm for 5 min (Centrifuge 5424, FA-45-24-11, Eppendorf). Recombinant *E. coli* cells were grown overnight in 5 mL selective liquid LB at 37°C and 150 rpm. 1.5-3 mL cell suspension were harvested in 1.5 mL reaction tubes. The supernatant was discarded, and the pellet was resuspended by vortexing in 150 µl P1-buffer (50 mM Tris, 10 mM EDTA, 100 µg/mL RNAseA, pH 8.0 with HCl). Cells were lysed in 150 µl P2-buffer (200 mM NaOH, 1 vol-% SDS) and inverted five times. Proteins were precipitated by adding 250 µl P3-buffer (2.55 M Na-acetate, pH 4.8 with acetic acid). The samples were inverted five times and centrifuged. Subsequently, 500 µL of the supernatant were transferred into new 1.5 mL tubes and mixed with 500 µl isopropanol. The samples were quickly vortexed and centrifuged again. Afterwards, the supernatant was discarded. At this step, the precipitated and non-visible plasmid-DNA pellet remained on the bottom of the tube. The pellet was washed twice with ice-cold EtOH (70 vol-%) omitting resuspending the DNA. After the second washing, the supernatant was discarded completely and the remaining EtOH was first removed by snapping the tube on a piece of clean paper towel and then through drying at 50-65°C for 10 min. The dried pellet was resuspended in 30 µL elution buffer (Tris/EDTA, pH 7.2) or deionized water. Purified plasmid-DNA with a concentration of 250-500 ng/µL was clean enough for subsequent cloning steps and test-digestion, however, an additional clean-up with the QIAquick PCR Purification Kit (Qiagen) was carried out when a Sanger sequencing reaction was necessary. P1-buffer needed to be stored at 4°C to maintain RNAse activity for up to 3 months. P2- and P3-buffer were stored at room temperature.

### A modified conjugation protocol for *C. ljungdahlii*

This protocol was adapted and modified according to Mock *et al.* (40). Cells of *C. ljungdahlii* were grown in RCM overnight to mid exponential growth until an OD_600_ of 0.4-0.8 was reached (NanoPhotometer® NP80, Implen). *E. coli* HB101 pKR2013 (DSM 5599) harboring the desired CRISPR-Cas12a-plasmid was grown as pre-culture in selective 5 mL LB medium overnight. The plasmid pKR2013 contains essential genes to mediate conjugation and a kanamycin resistance cassette. 1-2 mL of the *E. coli* cells were used to inoculate 10 mL selective LB medium in 50 mL baffled flask and cultivated until mid-exponential growth (OD_600_ 0.5-1.0). Subsequently, the *E. coli* culture was cooled to 4°C and 2 mL were transferred into sterile 2 mL reaction tubes. The *C. ljungdahlii* culture was kept at room temperature until use. Inside the anaerobic chamber, *E. coli* cells were centrifuged softly at 2900 rpm (mySpin^TM^ 12 mini centrifuge, Thermo Fischer Scientific) to protect pili, and washed once with sterile and anaerobic 0.1 M phosphate buffer saline (PBS) at pH 6.0. Afterwards, the washed pellet was resuspended gently in 100-150 µL cell suspension of *C. ljungdahlii* and directly transferred to well-dried RCM-agar plates (2 vol-% agar). Spot-mating was carried out at 37°C inside the anaerobic chamber overnight. After 8-24 h the spot was resuspended with anaerobic PBS (pH 6.0) and centrifuged at 10000 rpm (mySpin^TM^ 12 mini centrifuge, Thermo Fischer Scientific) for 3 min. The supernatant was discarded, and the pellet was resuspended in the remaining volume of the tube. Subsequently, 100 µL of the cell suspension was plated onto selective PETC+5gS-agar plates, which contained 5 g/L of peptone and 5 g/L meat extract to support growth. Selective agar plates should not be older than 2-3 days. Thiamphenicol was added for plasmid selectivity. Trimethoprim (10 mg/mL) was added to counter-select against *E. coli*. Growth was obtained after 4-5 days at 37°C inside the anaerobic chamber. *C. ljungdahlii* colonies were transferred into Hungate tubes containing 5 mL RCM with the respective antibiotics. A successful transformation of *C. ljungdahlii* with the correct plasmid was confirmed as follow: *1)* Growth in selective RCM with a characteristic pH decrease due to acetogenesis; *2)* control PCRs with primers for plasmid specific fragments; and *3)* plasmid purification from the culture and re-transformation into *E. coli* TOP10 cells.

### Electroporation of *C. ljungdahlii* cells

Electroporation of *C. ljungdahlii* cells was performed as previously reported (Xia et al. 2020) and applied for all non-CRISPR-based plasmids. Single colonies growing on selective plates were verified by PCR analyses and by re-transformation of plasmid DNA, which was extracted from *C. ljungdahlii* into *E. coli*.

### Growth experiments with *C. ljungdahlii*

In general, all recombinant *C. ljungdahlii* strains were pre-grown in 50 mL RCM in 100 mL serum bottles for 24-48 h. Subsequently, 2 mL cell suspension were used to inoculate 50 mL PETC medium in 100 mL serum bottles. This PETC pre-culture was cultivated for 40-48h at 37°C until mid-exponential growth phase at OD_600_ of 0.5-1.0. Afterwards, cells were transferred anaerobically into 50 mL reaction tubes, which were equilibrated for 3-5 days inside the anaerobic chamber. Cell harvest was performed outside the anaerobic chamber at 3700 rpm for 12 min (Centrifuge 5920R, S-4×1000, Eppendorf) at room temperature. After the centrifugation, the tubes were transferred back immediately into the anaerobic chamber, to keep the time at aerobic conditions at a minimum. Inside the anaerobic chamber, the supernatant was discarded, and the pellet was resuspended in fresh PETC medium to adjust to an OD_600_ of 5-10. The concentrated cell suspension was then transferred into sterile and anaerobic 10 mL Hungate tubes, sealed carefully, and used to inoculate main cultures outside of the anaerobic chamber. 1 mL of the cell suspension was used to inoculate 100 mL PETC main cultures. For heterotrophic growth experiment, 240 mL serum bottles were used. Autotrophic growth experiments were performed in 1000 mL Duran pressure plus bottles (Schott), to provide a high medium-to-headspace ratio. The Duran pressure plus bottles were sealed with butyl stoppers and a GL45 ring cap. Before inoculation of autotrophic cultures, the N_2_ headspace was replaced with a sterile gas mixture consisting of H_2_/CO_2_ (80/20 vol-%). Each bottle contained 0.5 bar overpressure. All cultures were cultivated in biological triplicates as batch cultures. The gas headspace was not refilled during the experiments. However, for the strain *C. ljungdahlii* pMTL83151_P_nat__*rnfCDGEAB* and the control strain *C. ljungdahlii* pMTL83151 we refilled the headspace during this experiment with the same gas mixture to 0.5 bar overpressure at time points 44.5 h, 73.5 h, and 148.5 h (**Supplementary Fig. S3**). Culture samples of 3 mL were taken at the bench and used for: *1)* OD_600_ measurement; *2)* pH measurement; *3)* HPLC analyses (acetate and ethanol); and *4)* FIA analyses (nitrate, nitrite, and ammonium). All culture samples were stored at −20°C until use. OD_600_ samples were diluted with medium or PBS buffer when the absorbance was > 0.5. We applied a two-tailed student’s t-test for all cultivation data. All p-values (*P*) below 0.001 indicate high significance and are given as ≤ 0.001.

### HPLC analyses

HPLC analyzes were performed as described before (24). In addition, all frozen supernatant samples were thawed at 30°C for 10 min and 300 rpm, vortexed briefly, and centrifuged for 3 min at 13806 rpm (Centrifuge 5424, FA-45-24-11, Eppendorf) before use. All HPLC samples were randomized.

### Measurement of nitrate, nitrite, and ammonium

Nitrate and nitrite concentrations were measured in a FIA continuous-flow analyzer system (AA3 HR AutoAnalyzer System, Seal Analytical GmbH, Germany) as described before (41). Briefly, nitrate is reduced to nitrite with hydrazine and then reacts with sulfanilamide and NEDD (N-1-Naphthylethylenediamine di-HCl) to form a pink complex, which can be quantified photometrically at 550 nm. The protocol follows the DIN 38405/ISO 13395 standard methods. Ammonium concentrations were measured in the same system but with salicylate and dichloroisocyanuric acid forming a blue complex that is measured at 660 nm instead. The protocol was following DIN 38406/ISO 11732 standard methods. Culture samples of *C. ljungdahlii* were treated as explained above for HPLC preparation. However, we prepared 1:50 dilution in 1 mL with deionized water *prior* to the FIA analyses. Standards for nitrate, nitrite, and ammonium were measured before and during the analyses for a standard curve and to minimize drift effects. Nitrate concentrations of each sample were calculated by the difference of the amount of nitrite measured with and without the *prior* reduction by hydrazine.

### Growth experiment for RNA extraction from *C. ljungdahlii*

For the expression analyses, we grew the strains *C. ljungdahlii* WT, *C. ljungdahlii* ΔRNF, and *C. ljungdahlii* Δ*rseC* under autotrophic and heterotrophic conditions as described above. The cultivation medium was PETC with ammonium as nitrogen source. Pre-cultures were grown in heterotrophic medium for 48 h. Next, the cells were transferred into the anaerobic chamber and harvested for 12 min at 25°C and 3700 rpm (Centrifuge 5920 R, S-4×1000, Eppendorf) outside of the anaerobic chamber. The supernatant was discarded under anaerobic conditions and the pellet was resuspended in fresh medium of the main cultures. The start OD_600_ for autotrophic main cultures was 0.2, while it was 0.15 for heterotrophic conditions. The cultures were cultivated at 37°C. 10 mL culture samples were taken after 3 h and 20 h. The samples were immediately cooled on ice and centrifuged for 12 min at 4°C and 3700 rpm (Centrifuge 5920 R, S-4×1000, Eppendorf). The cell pellets were stored at −20°C until RNA extraction.

RNA was purified from *C. ljungdahlii* with the RNeasy Mini Kit (Qiagen) as described before (42). For the RNA extraction, we used 2·10^8^ cells, which was approximately 10 mL of a *C. ljungdahlii* culture at OD_600_ 0.2. The cell lysis was performed in the lysis buffer of the kit with 50 mg glass beads (0.1 mm silica spheres, MP Biomedicals) in a bead beater (5G-FastPrep, MP Biomedicals) for 2x 60s at 9 m/s. RNA samples were eluted in 30 µL nuclease-free water. After the extraction procedure, an additional DNase I digest (RNase free Kit, Thermo-Scientific) was performed to remove potential DNA contamination. Elimination of genomic DNA was confirmed with PCR analyses and gel electrophoresis. cDNA synthesis was performed with the QuantiTect Reverse Transcriptase Kit (Qiagen) according to the manufactureŕs instructions. We used 500 ng RNA as template for each reaction. cDNA was stored at −20°C until further use.

### qRT-PCR analyses

All qRT-PCR analyses were performed in a Quantstudio 3 Thermocycler (Applied Biosystems, Thermo Scientific). The PCR reaction mix contained 10 µL SYBR Green Master mix (Thermo Scientific), 1 µL of a fwd and rv qRT-PCR primer (final concentration 500 nM) (Table 5), and 1 µL (∼5 ng) cDNA template. We used the *rho* gene as reference gene, which was described before as suitable candidate for qRT-PCR experiments with *C. ljungdahlii* (42). We added RNA controls to further exclude gDNA contamination in our samples. All qRT-PCR reactions were performed in technical triplicate according to the manufacturer’s instructions. We set the Ct threshold to 0.1. The fold change in gene expression between the samples was determined with the 2^-ΔΔCt^ method as described before (43). We examined the PCR efficiency of our qPCR master mix by using plasmid DNA containing the sequence of each tested gene in a series of dilutions (10^-1^, 10^-2^, 5·10^-3^, 10^-3^, 5·10^-4^, 10^-4^). The slopes were ranging from 0.04-0.09 for the RNF-gene cluster genes and 0.17 for *rseC*, and were thus, close to zero, which proofs that the efficiencies are similar and the 2^-ΔΔCt^ can be used for interpretation of the qRT-PCR data (43). We applied a two-tailed student’s t-test based on our ΔCt values for each gene to analyze the significance of our samples in comparison to the wild type.

### Strain preservation

Cultures of *C. ljungdahlii* were stored at −80°C. For this, cultures were grown in RCM until late exponential growth phase (OD_600_ 0.8-1.2) at 37°C for 36-48 h. The cells were transferred into anaerobic 50 mL reaction tubes inside the anaerobic chamber and harvested outside of the anaerobic chamber for 12 min at 4°C and 3700 rpm (Centrifuge 5920 R, S-4×1000, Eppendorf). The supernatant was discarded inside the anaerobic chamber and the pellet was resuspended in fresh RCM medium to an OD_600_ of 5-10. 2 mL of the cell suspension was transferred into 10 mL serum bottles, which were previously filled with 2 mL of 25-50 vol-% anaerobic and autoclaved glycerol. The serum bottles were briefly vortexed outside the anaerobic chamber, incubated on ice for 10-15 min and subsequently frozen at −80°C. For inoculation of a new RCM culture, a single serum bottle was quickly thawed up under rinsing water and 1-2 mL of the cell suspension was immediately transferred with a syringe into the medium bottle. Cultures of *E. coli* were stored at-80°C in sterile screw-cap tubes filled with 25-50 vol-% glycerol.

## Acknowledgement

This work was funded through the Alexander von Humboldt Foundation in the framework of the Alexander von Humboldt Professorship, which was awarded to L.T.A. We are also thankful for additional funding to L.T.A. and B.M. from the Deutsche Forschungsgemeinschaft (DFG, German Research Foundation) under Germany’s Excellence Strategy – EXC 2124 – 390838134. The plasmid pMTL2tet01gusA was kindly provided by Dr. Gregory Stephanopoulos (Department of Chemical Engineering, Massachusetts Institute of Technology). We acknowledge support by the DFG and Open Access Publishing Fund of the University of Tübingen. We thank Dr. Peng-Fei Xia (Environmental Biotechnology Group, University of Tübingen) for his advice for the CRISPR design and Franziska Schädler (Geomicrobiology and Microbial Ecology, University of Tübingen) for her support with the nitrate and ammonium measurements. We thank Nicole Smith (Environmental Biotechnology Group, University of Tübingen) for her support in medium preparations.

## Author’s contribution

Christian-Marco Klask (C.M.K.) and Bastian Molitor (B.M.) designed the experiments. C.M.K. performed the genetic work, conducted the growth experiments, analyzed the metabolites, and performed the *in-silico* research. C.M.K. and Benedikt Jäger (B.J.) performed the qRT-PCR experiments. C.M.K. analyzed the experimental data. Largus T. Angenent (L.T.A.) and B.M. supervised the work. C.M.K. and B.M. wrote the manuscript, and all authors edited the paper and approved the final version.

## Supplementary Figure legends

**Supplementary Fig. S1 Heterotrophic growth and metabolic products of C. ljungdahlii WT, ΔRNF, and ΔrseC.** Cultures of *C. ljungdahlii* WT (●, ○), ΔRNF (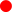, 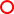), and Δ*rseC* (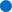, 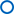) were grown in 100 mL PETC medium in 240 mL bottles at 37°C. The headspace consisted of N_2_ (100 vol-%). Fructose (5 g/L) was added as carbon source. The medium contained either 18.7 mM nitrate (NO_3_^-^) (filled circles) or 18.7 mM ammonium (NH_4_^+^) (open circles) as nitrogen source. The cultivation times were 79 h for the WT and ΔRNF strain, and 84 h for the Δ*rseC* strain. All cultures were grown in biological triplicates, data is given as mean values, with error bars indicating the standard deviation. **A**, growth; **B**, pH-behavior; **C**, acetate concentrations; **D**, ethanol concentration; **E**, ammonium concentration; and **F**, nitrate concentrations. WT, wild type; ΔRNF, RNF-gene cluster deletion; Δ*rseC*, *rseC* gene deletion.

**Supplementary Fig. S2 CRISPR-Cas12a-mediated *rseC* gene and nar gene cluster deletion in C. ljungdahlii. A,** verification of the *rseC* gene deletion. PCR-samples for the *fdhA* fragment (WT: 501 bp, deletion strain: 501 bp), *rseC* fragment (WT: 417 bp, deletion strain: no fragment), and for a fragment that was amplified with primers that bind 1104 bp upstream and 1208 bp downstream of the *rseC* gene locus (WT: 2755 bp, deletion strain: 2338 bp). DNA-template: gDNA of *C. ljungdahlii* Δ*rseC* (lane A1, A4, and A7); gDNA of *C. ljungdahlii* WT (lane A3, A6, and A9); and water (lane A2, A5, A8). **B**, verification of the *nar* gene cluster deletion PCR samples for the *fdhA* fragment (WT: 501 bp, deletion strain: 501 bp), *nar* fragment (WT: 3739 bp, deletion strain: no fragment), and for a fragment that was amplified with primers that bind 1137 bp upstream and 1110 bp downstream of the *nar* gene cluster locus (WT: 5986 bp, deletion strain: 2247 bp). DNA-template: gDNA of *C. ljungdahlii* Δ*nar* (lane B1, B4, and B7); gDNA of *C. ljungdahlii* WT (lane B3, B6, and B9); and water (lane B2, B5, B8). M: Generuler^TM^ 1 kb.

**Supplementary Fig. S3 Autotrophic growth and metabolic products of the overexpression strains *C. ljungdahlii* pMTL83151_Pnat_*rnfCDGEAB* and *C. ljungdahlii* pMTL83152_*rseC*.** Cultures were grown in 100 mL PETC medium in 1 L bottles at 37°C and 150 rpm. The headspace consisted of H_2_ and CO_2_ (80/20 vol-%) and was set to 0.5 bar overpressure. For the strain *C. ljungdahlii* pMTL83151_P_nat__*rnfCDGEAB* and the control strain *C. ljungdahlii* pMTL83151 we refilled the headspace during this experiment with the same gas mixture to 0.5 bar overpressure at time points 44.5 h, 73.5 h, and 148.5 h. The medium contained 18.7 mM ammonium as nitrogen source. Thiamphenicol (5 µg/mL) was used for selection. All cultures were grown in biological triplicates, data is given as mean values, with error bars indicating the standard deviation. The cultivation time was 185 h and 197 h for *C. ljungdahlii* pMTL83151_P_nat__*rnfCDGEAB* and *C. ljungdahlii* pMTL83152_*rseC*, respectively. (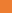) *C. ljungdahlii* pMTL83151_P_nat__*rnfCDGEAB*; (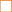) *C. ljungdahlii* pMTL83151 (empty plasmid); (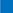) *C. ljungdahlii* pMTL83152_*rseC*; (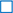) *C. ljungdahlii* pMTL83152 (empty plasmid). *A*, growth; *B*, pH-behavior; *C*, acetate concentrations; and *D*, ethanol concentration. rpm, revolutions per minute; CO_2_, carbon dioxide; and H_2_, hydrogen.

**Supplementary Fig. S4 Gene expression change of the *rnfCDGEAB* cluster genes and the *rseC* gene in the wild-type strain from heterotrophy to autotrophy. A,** gene expression change after 3 h cultivation time; **B**, gene expression change after 20 h cultivation time. RNA samples were purified from cultures that were cultivated either autotrophically with hydrogen and carbon dioxide or heterotrophically with fructose. cDNA was synthesized from the purified RNA samples and used as template for qRT-PCR analyses. The *rho* gene was used as “housekeeping” gene. The fold change in gene expression was determined with the 2^-ΔΔCT^ method (43). ***, *P* ≤ 0.001; and *ns, not significant (*P* > 0.5). We defined log_2_ (fc) ≤ −1 as downregulated genes and ≥ +1 as upregulated genes.

**Supplementary Fig. S5 Multiple sequence alignment of RseC amino-acid sequence using CLUSTAL Omega.** The symbols indicate low similarity (.), high similarity (:), and identical amino acids (*) between the amino acid sequences. Similar colors indicate similar amino acids. The type strains were *C. ljungdahlii* DSM13528; *C. autoethanogenum* DSM10061; *C. carboxidovorans* P7; *C. kluyveri* DSM555; *E. limosum* ATCC8486; *A. woodii* DSM1030; *R. capsulatus* SB1003; and *E. coli* K-12. Clustal omega version 1.2.4. with default settings was used for the analysis (https://www.ebi.ac.uk/Tools/msa/clustalo/, 05/2021).

**Supplementary Fig. S6 Heterotrophic growth and metabolic products of *C. ljungdahlii* Δnar.** Cultures were grown in 100 mL PETC medium in 240 mL bottles at 37°C. Fructose (5 g/L) was added as carbon source. The headspace consisted of N_2_ (100 vol-%). The medium contained either 18.7 mM nitrate (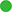) or 18.7 mM ammonium (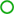) as nitrogen source. All cultures were grown in biological triplicates, data is given as mean values, with error bars indicating the standard deviation. The cultivation times was 84 h. The *C. ljungdahlii* WT data (●, ○) from Supplementary Fig. S1 is given for comparison. **A**, growth; **B**, pH-behavior; **C**, acetate concentrations; **D**, ethanol concentration; **E**, ammonium concentration; *F*, nitrate concentrations; and *G*, fructose concentrations. Δ*nar*, deletion of the nitrate reductase genes.

**Supplementary Fig. S7 Autotrophic growth and metabolic products of plasmid-based complemented strain *C. ljungdahlii Δnar* pMTL83152_*nar*.** Cultures were grown in 100 mL PETC medium in 1 L bottles at 37°C and 150 rpm. The headspace consisted of H_2_ and CO_2_ (80/20 vol-%) and was set to 0.5 bar overpressure. The medium contained 18.7 mM nitrate (NO_3_^--^) but no ammonium (NH_4_^+^) as nitrogen source. Thiamphenicol (5 µg/mL) was used for selection. All cultures were grown in biological triplicates, data is given as mean values, with error bars indicating the standard deviation. The cultivation times was 192.5 h. (●) *C. ljungdahlii* Δ*nar* pMTL83152_*nar*; (○) *C. ljungdahlii* Δ*nar* pMTL83152 (empty plasmid); **A**, growth, **B**, pH-behavior; **C**, acetate concentrations; **D**, ethanol concentration; **E**, ammonium concentration; and **F**, nitrate concentrations. Δ*nar*, gene deletion of the nitrate reductase genes; rpm, revolutions per minute; CO_2_, carbon dioxide; and H_2_, hydrogen.

